# MorphoGraphX 2.0: Providing context for biological image analysis with positional information

**DOI:** 10.1101/2021.08.12.456042

**Authors:** Soeren Strauss, Adam Runions, Brendan Lane, Dennis Eschweiler, Namrata Bajpai, Nicola Trozzi, Anne-Lise Routier-Kierzkowska, Saiko Yoshida, Sylvia Rodrigues da Silveira, Athul Vijayan, Rachele Tofanelli, Mateusz Majda, Emillie Echevin, Constance Le Gloanec, Hana Bertrand-Rakusova, Milad Adibi, Kay Schneitz, George Bassel, Daniel Kierzkowski, Johannes Stegmaier, Miltos Tsiantis, Richard S. Smith

## Abstract

Positional information is a central concept in developmental biology. In developing organs, positional information can be idealized as a local coordinate system that arises from morphogen gradients controlled by organizers at key locations. This offers a plausible mechanism for the integration of the molecular networks operating in individual cells into the spatially-coordinated multicellular responses necessary for the organization of emergent forms. Understanding how positional cues guide morphogenesis requires the quantification of gene expression and growth dynamics in the context of their underlying coordinate systems. Here we present recent advances in the MorphoGraphX software (Barbier de Reuille et al. eLife 2015;4:e05864) that implement a generalized framework to annotate developing organs with local coordinate systems. These coordinate systems introduce an organ-centric spatial context to microscopy data, allowing gene expression and growth to be quantified and compared in the context of the positional information thought to control them.

## Introduction

Many aspect of animal morphogenesis are thought to be controlled by positional information (Wolpert, 1969), where cells can sense their position in a developing organ and respond accordingly. This phenomenon may be even more pervasive in plants, as cells cannot relocate within organs, and must decide their fate based on their location. For example, root morphogenesis appears to be controlled by an organizing center at the root tip that provides founder cells and positional information to the growing structure (Scheres et al., 2002). Ablation of cortical cell initials in the root meristem causes the neighboring pericycle cells to divide and fill the available space, subsequently adopting the fate associated to their new location (van den Berg et al., 1995). A similar effect his has been demonstrated for a variety of cell types in the Arabidopsis root (Marhava et al., 2019). In leaves, development is thought to be coordinated by polarity fields oriented from leaf base to tip (Kierzkowski et al., 2019; Kuchen et al., 2012). Over time organs can initiate new growth axes, such as when serrations or leaflets develop in more complex leaves (Barkoulas et al., 2008; Kierzkowski et al., 2019), or lateral roots emerge from the primary root (Scheres et al., 2002). In these cases information from several organizers must be integrated to direct cell response.

To understand how positional information controls morphogenesis, it is necessary to quantify cell shape, gene expression and morphogen concentration changes over time, preferably at the cellular level. This information then needs to be related to its position relative to the organizers controlling development within the organ. As computational power and imaging methods improve, new software packages for cell segmentation and lineage tracking are being developed (Sommer et al., 2011; Stegmaier et al., 2016), including many specialized for plants (Barbier de Reuille et al., 2015; Eschweiler et al., 2019; Fernandez et al., 2010; Schmidt et al., 2014; Wolny et al., 2020). This progress has enabled the segmentation of time-lapse data at increasingly higher resolution and throughput (Hervieux et al., 2016; Kierzkowski et al., 2019; Sapala et al., 2018; Willis et al., 2016)). Although this increase in data volume offers tremendous potential to understand how genes control form, the analysis of geometric data from thousands of cells is non trivial. Information about a cell’s shape, gene expression and growth directions is of limited value when the cell’s spatial context within the developing organ is unknown.

MorphoGraphX is a computer software platform that is specialized for image processing on surface layers of cells (Barbier de Reuille et al., 2015). It has proven especially useful for the analysis of confocal microscopy images from time-lapse data in order to quantify the cellular level dynamics of growth, cell division and gene expression (e.g. Bringmann & Bergmann, 2017; Feng et al., 2018; Hervieux et al., 2016; Hong et al., 2016; Kierzkowski et al., 2019; Louveaux et al., 2016; Sapala et al., 2018; Scheuring et al., 2016; Tsugawa et al., 2017; Vlad et al., 2014; Zhang et al., 2020; Zhu et al., 2020). Key to the approach taken in the software is the representation of cell layers as curved, triangulated surface meshes that capture the overall 3D shape of organs, which retains much of the simplicity of 2D segmentation and lineage tracking. These “2.5D” images contain the geometry of the sample at two scales. The global shape of the organ is captured by the mesh’s geometry, while a cellular-scale representation is obtained from the confocal signal projected onto the mesh, which is segmented to extract the shape of individual cells on the surface (Fig. 1A-C). When combined with time-lapse data acquisition and cell lineage tracking, MorphoGraphX allows cell growth and its relationship to gene expression to be quantified calculated (Fig. 1D,E; Kierzkowski et al., 2019; Sapala et al., 2018; Vlad et al., 2014). In addition to cell surface analysis, MorphoGraphX also supports the creation and analysis of full 3D meshes with volumetric cells (Fig. 1S1, Vijayan et al., 2021). Here we describe new methods we have developed in MorphoGraphX to understand these data by additionally annotating cells with positional information. Not unlike the annotation of sequence data, this allows cellular data to be given spatial context, and a frame of reference within the organ relative to its developmental axes and the organizers instructing morphogenesis.

**Figure 1:**
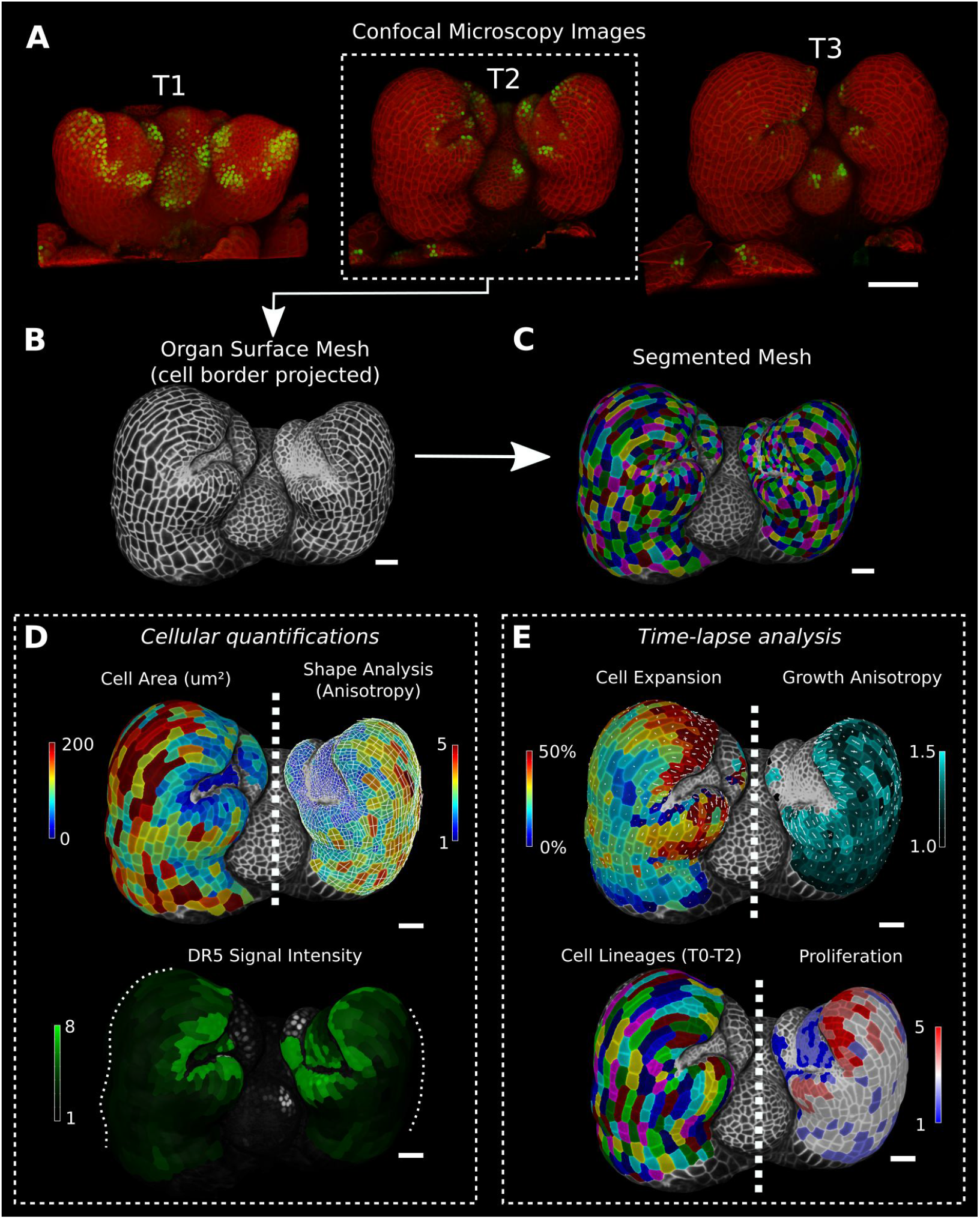
Cellular segmentation and basic quantifications supported by MorphoGraphX demonstrated by using a time-lapse series of an *A. thaliana* flower meristem using MorphoGraphX. (A) Multi-channel confocal microscopy images with a cell wall signal (red) and DR5 marker signal (green). Shown are the last 3 time points (T1-T3) of a 4-image series (T0-T3). (B-C) Extracted surface mesh of T2. Cell wall signal near the surface was projected onto the curved mesh to enable the creation of the cellular segmentation in (C). The segmented meshes provide the base for further analysis within MorphoGraphX as shown in (D) and (E). (D) Top: MorphoGraphX allows the quantification of cellular properties such as cell area and shape anisotropy (shown as heat maps). The white axes show the max and min axes of the cells. Bottom: Heat map of the quantification of the DR5 marker signal (arbitrary units) projected onto the cell surface mesh. (E) When cell lineages are known, time-lapse data can be analyzed. Top: Heat maps of areal growth and growth anisotropy (computed from T1 to T2). The white lines inside the cells depict the principal directions of growth. Bottom: Visualization of the cell lineages from T0 to T2 and a heat map of cellular proliferation (number of daughter cells). Scale bars: (A) 50μm; (B - E) 20μm.

**Figure 1S1:**
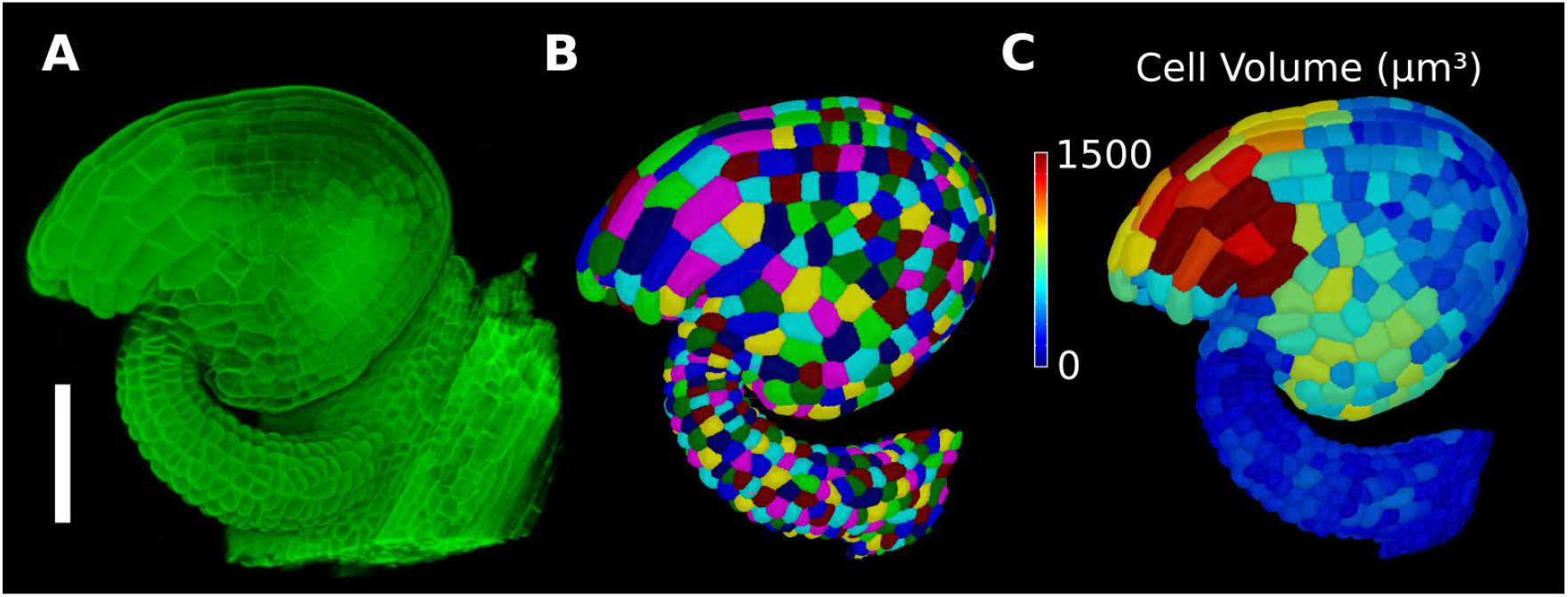
Basic 3D analysis using MorphoGraphX demonstrated using an Arabidopsis ovule. (A) Confocal microscopy image with cell wall staining. (B) Segmented mesh with volumetric cells. (C) The segmented mesh allows cellular geometry to be quantified. Shown is a heat map of cell volumes. Scale bar: 50μm.

## Results

### Defining directions within an organ

The simplest method to provide positional information for the cells in a sample is by aligning the sample with a set of 3D coordinate axes (Fig 2A). For example, a developing root meristem can be aligned and positioned such that the organizing quiescent center is at the origin with the Y-axis increasing in the longitudinal direction of the root. Provided the sample is reasonably straight, this allows cellular measures to be compared with their distance from the quiescent center (Fig 2B) (Schmidt et al., 2014).

**Figure 2:**
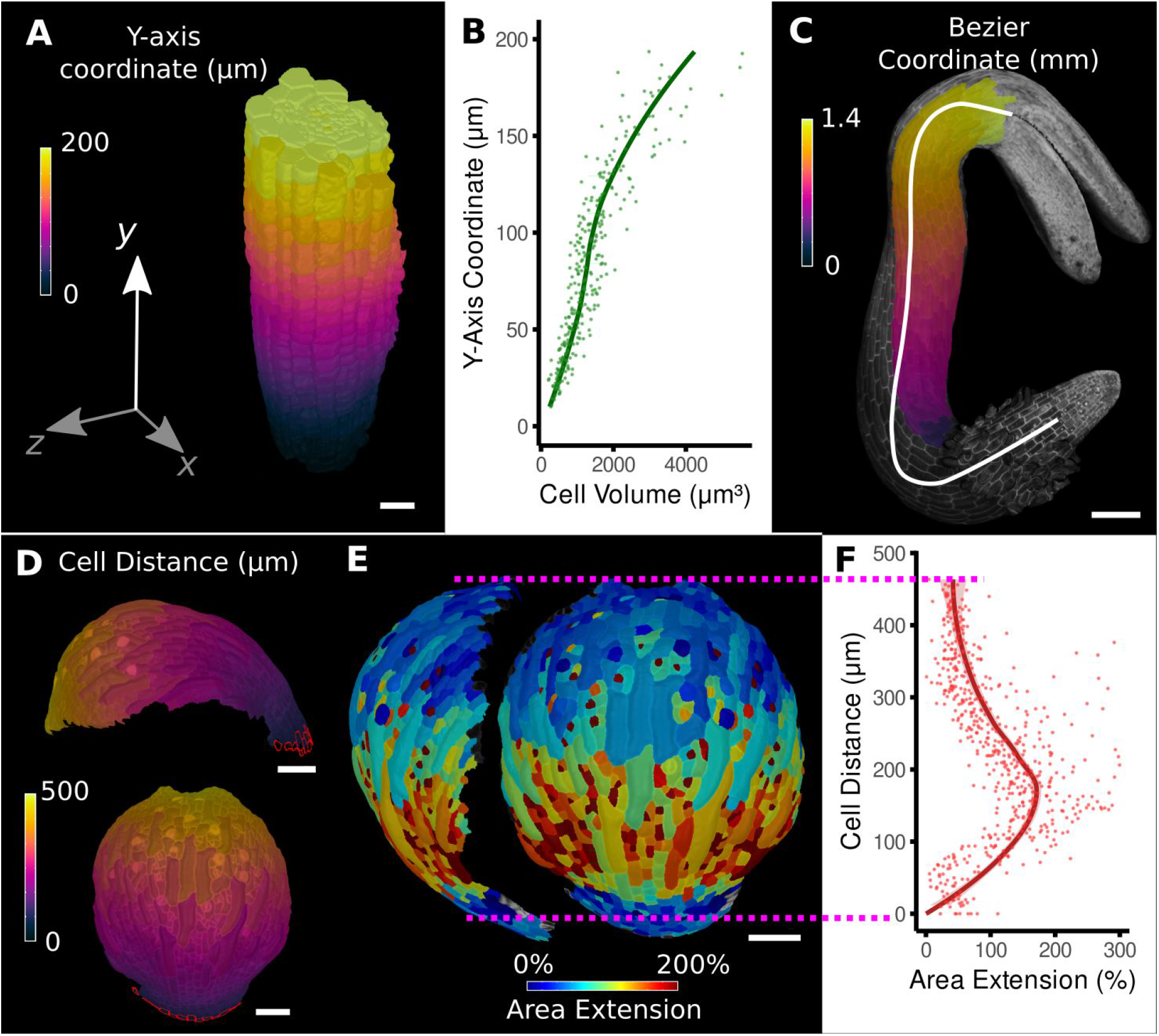
Methods to define positional information and their application to data analysis in plant organs. (A) Y-axis aligned *A. thaliana* root. The cells are colored according to the y-coordinate of their centroid position. (B) Plot of cell volumes of epidermis cells of the root in (A) along the y-axis with a fitted trend line. (C) Seedling of *A. thaliana* with a surface segmentation of the epidermis. A manually defined Bezier curve (white) allows the assignment of accurate cell coordinates along a curved organ axis. (D) Side and top view of an *A. thaliana* sepal with a proximal-distal (PD) axis heat coloring. The cell coordinates were assigned by computing the distance to manually selected cells (outlined in red) at the organ base. This method allows organ coordinates to be assigned in highly curved tissues. (E) Side and top view of (D) with a heat map coloring based on cellular growth to the next time point. (F) Plot summarizing the growth data of (E) using the PD-axis coordinates from (D). See Figure 2S1 for the analysis of the complete time-lapse series. Scale bars: (A) 20 μm; (C) 100 μm; (D, E) 50 μm.

However, for highly curved organs significant errors will occur, especially in more distal regions, further from the origin. It is possible to overcome this problem by placing a curve along the central axis (Fig 2C, Montenegro-Johnson et al., 2015). For this curve, MorphoGraphX uses Bezier splines defined by control points. These points can be positioned to create an axis that conforms to the curvature of the organ, using either interactive manipulation of the control points, or automatically from a selected file of cells. Distance can then be calculated along the line, and transferred to cells in the cross section perpendicular to the line. MorphoGraphX also allows a 2D Bezier surface to be positioned next to or within a sample, enabling two directions to be aligned with the natural curvature of the sample.

An alternative method applicable to curved organs with more complex shape, is to select one or more cells at a reference position, and calculate the distance relative to the selection (Fig 2D, Movie 1). This offers an easy method to create a distance field, and greatly increases the variety of organs that can be accommodated. The distance is determined by computing the shortest path along cells through the tissue, causing it to naturally follow the curvature of the organ. As many organs have a layered cellular organization, a distance field normal to the sample’s surface can be calculated. The surface of the sample is extracted and is used to annotate a full 3D segmentation of the interior cells. Fig. 3A shows a shoot meristem with the cells colored by distance to the surface, which can be used to classify cells (Fig 3C).

**Figure 3:**
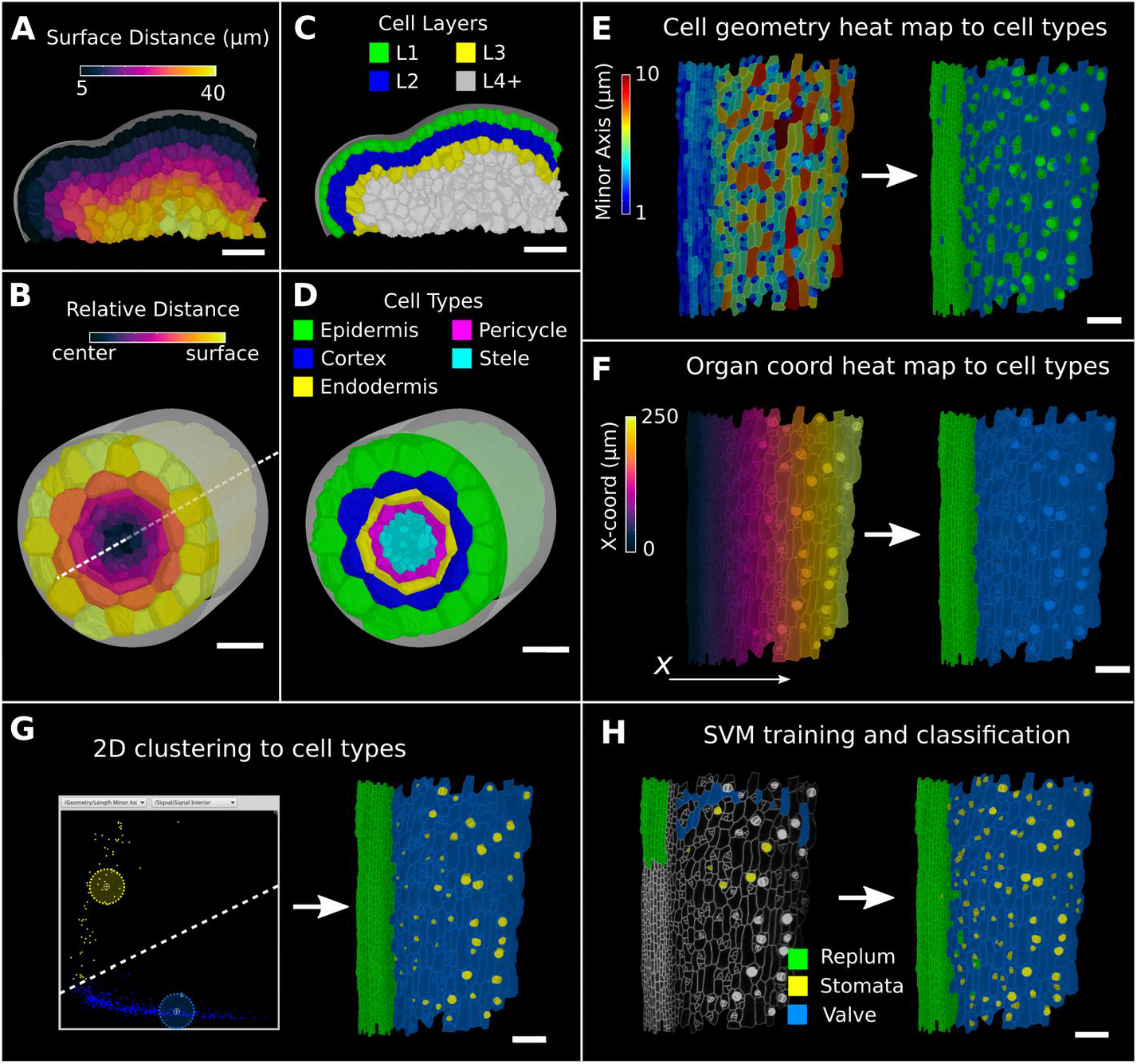
Methods to create organ coordinates for 3D meshes and to label different cell types. (A-D) Organ coordinates and cell types for volumetric meshes. (A) Heat map of the surface distance for cell centroids in an *A. thaliana* shoot apical meristem. (B) For volumetric tissues often a single direction is not enough to capture the geometry of the organ. Different methods can be combined such as a Bezier curve (white dashed line) with a surface mesh (grey) to create a heat map of the relative radial distance of cells in the *A. thaliana* root. (C-D) Organ coordinates can be used to assign cell type labels as demonstrated in the 3D Cell Atlas plugin for meristem and root. See also Figure 3S1. (E-H) Different methods to create cell type labellings. (E) *A. thaliana* gynoecium (fruit epidermis) surface segmentation with a heat map of the length of the minor cell axis as obtained from a PCA on the cells’ triangles. The heat values can be thresholded to assign two cell types. (F) The same principle can be used on organ coordinates which results in a clean separation of replum (green) and valve tissue (blue). (G) We generalized the 2D clustering approach of 3D Cell Atlas (see Figure 3S1) so that it can be used for any measure pair and on subset selections of cells. Shown is a 2D plot of the minor axis length (x-coord) and cell signal intensity (y-coord) on the valve tissue in (F). Manually assigning clusters can separate the stomata, which are typically smaller with higher signal values (yellow) and the remaining valve cells (blue) efficiently. See also Figure 3S1 for 2D plots of all cells. (H) The Support Vector Machine (SVM) classification is able to separate the 3 shown cell types in a higher dimensional space by using a variety of different measures and a relatively small training set. Scale bars: (A-D) 20 μm; (E-H) 50 μm.

**Figure 2S1:**
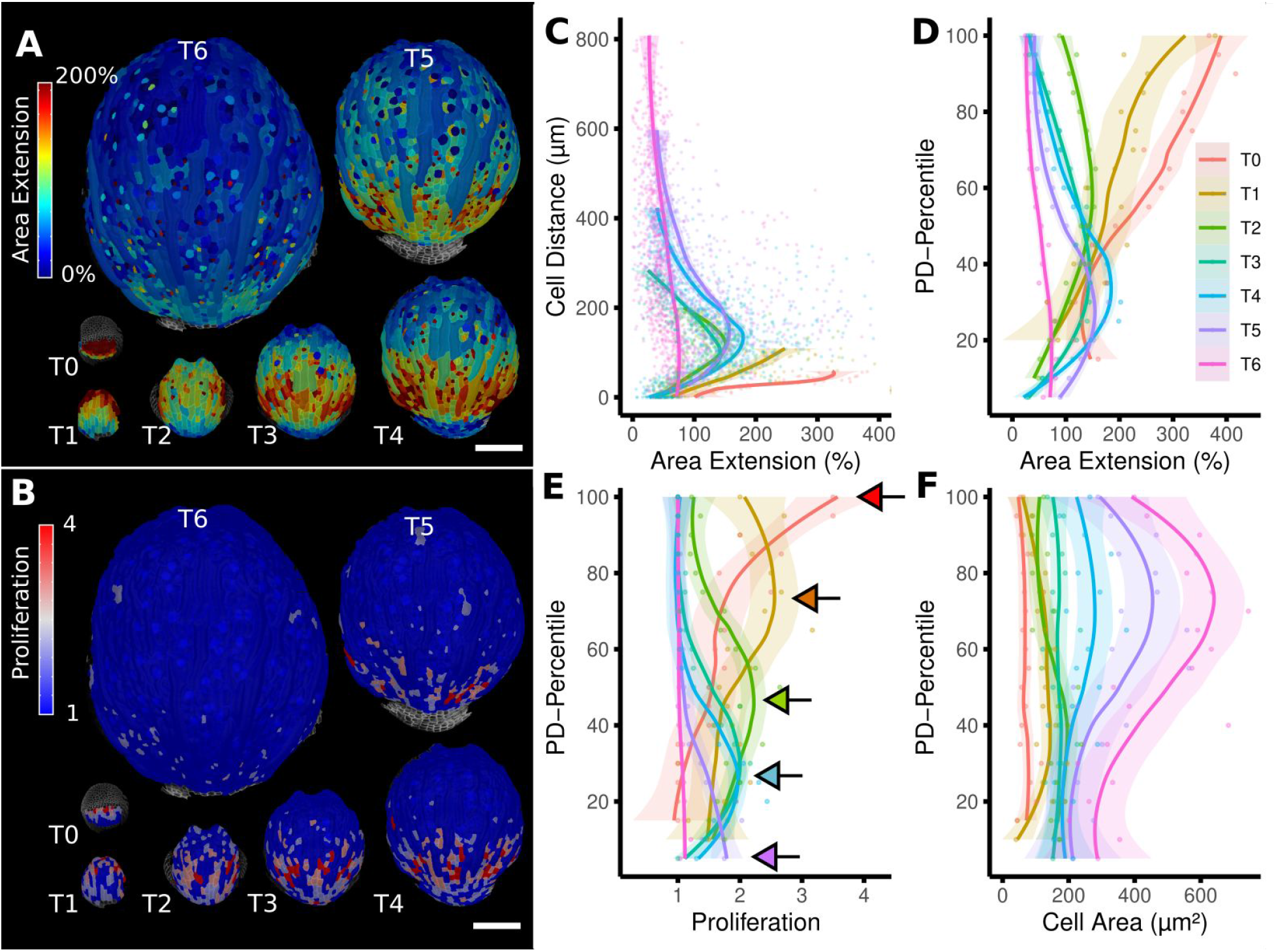
From cellular resolution heat maps to a global analysis of *A. thaliana* sepal development using organ-centric coordinates. (A-B) Heat maps of cell area extension (A) and cell proliferation (B) for each time point. (C) Plot of the heat map data from (A) vs the distance of cells to the base of the organ (see also Fig 2D). The distance of the maximum of the growth zone from the base or the organ is relatively constant. However, organ length is increases about 10x between the first and last time points, making a comparison of the different curves difficult. (D-F) When plotting the same data with normalized cell distance values averaged using 20 bins along the PD-axis it becomes more apparent that the growth zone moves from the proximal to the distal regions over the course of development (D). The trend of lower and more proximal maxima (highlighted with arrows) is even clearer when proliferation is plotted in the same way (E). (F) Cell area data plotted as in (D) and (E). Average cell areas increase mainly at the distal end during later time points. Scale bars: (A, B) 100 μm.

### Combining directions

In 2D or on 2.5D surfaces, local directions can be fully defined by a single distance measure, by taking one direction aligned with the gradient of the distance field or a Bezier curve, and the other perpendicular to the first. This is similar to methods used to specify directions in developmental modeling in plants (Green et al., 2010; Kennaway & Coen, 2019; Kierzkowski et al., 2019; Kuchen et al., 2012; Whitewoods et al., 2020), and thus facilitates direct comparison between models and experimentally observed patterns of growth and gene expression.

In 3D, a third direction must be defined (Kennaway & Coen, 2019; Whitewoods et al., 2020). In MorphoGraphX this can be done by combining the directions defined by different distance measures. The 3D Cell Atlas add-on for MorphoGraphX (Montenegro-Johnson et al., 2015) combines several distance measures for radially symmetric structures such as root and hypocotyls. A Bezier curve is placed along the center in the longitudinal direction and combined with a surface mesh to obtain radial directions (Figure 3B). This also puts bounds on the radial direction (and also implicitly on the circumferential direction) which allows relative coordinates to be assigned to cells in addition to absolute values. For less regular organs, like the ovule, coordinates derived from a surface mesh (Fig 4A) can be combined with a Bezier curve along the surface (Fig 4C) to define a coordinate system for the region of interest in full 3D. In both of these cases, relative coordinates facilitate the classification of cells into layers. When annotating a root using the 3D Cell Atlas coordinate system, the relative radial coordinate will follow the layer as the organ narrows towards the tip. The use of relative coordinates also makes it possible to pool data from multiple samples (Vijayan et al., 2021; Zhang et al., 2020) and to compare data from different genotypes (Kierzkowski et al., 2019; Montenegro-Johnson et al., 2019).

**Figure 4:**
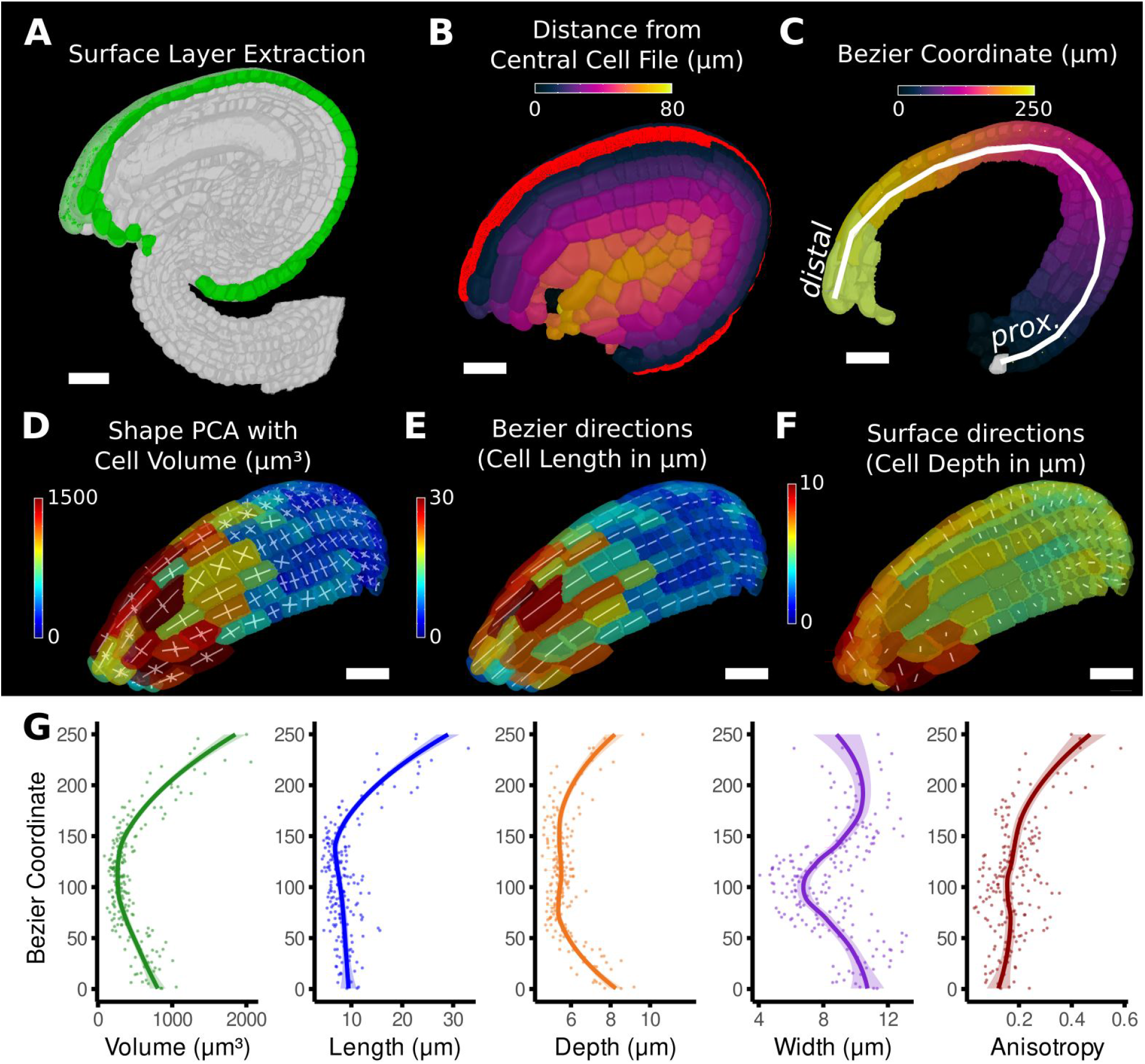
Quantification of volumetric cell sizes along organ axes in the outer layer of the outer integuement of an *A. thaliana* ovule. (A) Extraction of cell layer of interest (colored in green) using an organ surface mesh. (B) Selection of the central cell file (in red) with cell distance heat map to exclude lateral cells (heat values >40um). (C) The centroids of the selected cells from (B) were used to specify a Bezier curve defining the highly curved organ axis from the proximal to the distal side. Heat coloring of the cells according to their coordinate along the Bezier. (D-F) Analysis of the cellular geometry in 3D. (D) Heat map of cell volume and the tensor of the three principal cell axes obtained from a Principal Component Analysis on the segmented stack. (E) Bezier directions and associated cell length. (F) Directions perpendicular to the surface and associated cell depth. (G) Plots of the various cellular parameters relative to the Bezier coordinate. Scale bars: 20μm.

**Figure 3S1:**
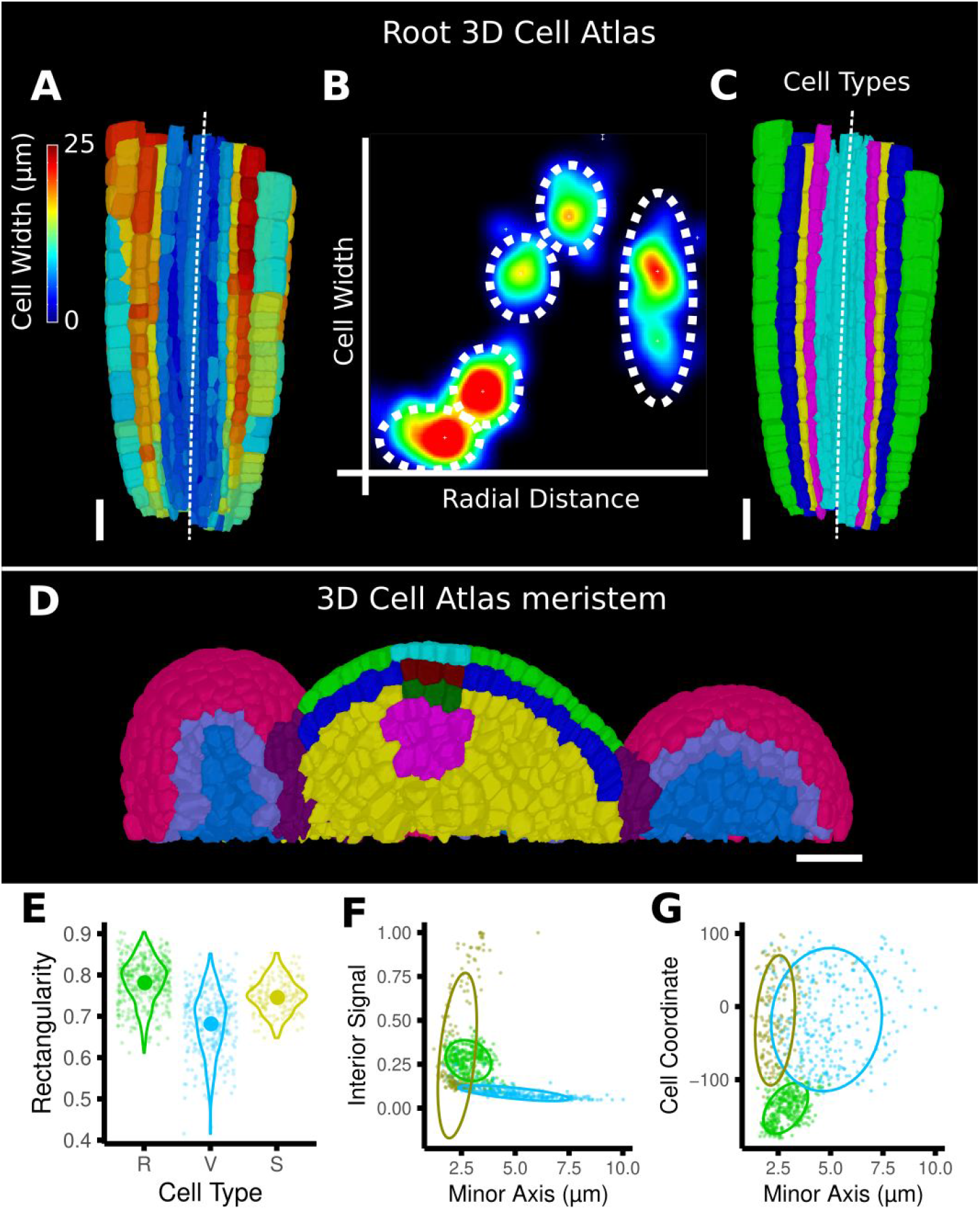
Cell type labeling methods and their use them in the data analysis. (A-C) Methods supported by the 3D Cell Atlas Add-on demonstrated on a *A. thaliana* root (Montenegro-Johnson et al., 2015). (A) Longitudinal cross section with a heat map of circumferential cell size. The white dashed line is the manually defined central axis. (B) The 2D heat plot of radial distance (heat map: in Fig 3B) and circumferential cell size (heat map in (A)) reveals a distinct clustering and can be used directly to assign the different cell types (dashed ellipses). (C) Final result of the cell type assignment. (D) The 3D cell atlas meristem (Montenegro-Johnson et al., 2019) allows the assignment of cell layers (as also seen in Figure 3C) and types in the meristem. (E-G) Cell type specific data analysis on the example of the *A. thaliana* gynoecium (Figure 3E-H). (E) Violin plot of the rectangularity of different cell types. Valve tissue cells are less rectangular and have higher variance compared to replum cells and stomata. (F-G) 2D scatter plots with fitting ellipses. Choosing different measures as x- or y-axis allows the separation of different cell types as demonstrated in Figure 3G. Scale bars: (A, C, D) 20μm.

### Using positional information for data analysis

Once cells have been annotated with positional information, it can be used to analyze cell-level data, such as growth, cell proliferation, cell shape and gene expression. Using the distance measure to define the proximal-distal axis (Fig 2D), geometric measures can be plotted against the local coordinate system. In Fig 2F cell area extension was plotted against distance from the base of the sepal. On the full 7 day sepal time-lapse shown in Fig 2S1 (Hervieux et al., 2016), it can be seen that initially cell growth is more distal, with a band of high growth progressing towards the base of the sepal. By time point 6, the growth has slowed and become more uniform as the organ differentiates. Proliferation is initially more uniform, but otherwise follows a pattern similar to growth, progressing basally as the organ matures. The data can be indexed by position and visualized in graphs, showing how growth and proliferation vary along a developmental axis (Fig. 2S1C-F). This also makes it possible to group data collected from multiple samples and to compare different genotypes (Kierzkowski et al., 2019; Zhang et al., 2020).

In addition to scalar information such as areal growth rate or cell volume, MorphoGraphX can also quantify directional information, such as the Principal Directions of Growth (PDGs) that represent the maximal and minimal directions of growth for each cell (Figure 5A-C). A common problem with the interpretation of such directional information is the tendency for directions to be locally heterogeneous when growth is nearly isotropic. This happens because the maximal and minimal growth amounts are almost the same, and the displayed directions become arbitrary, and heavily influenced by noise. This can make the comparison of growth directions between neighboring cells difficult. A more informative approach is to look at growth with respect to the directions of the developmental axes. This can be done by first setting up an axis defining the positional information for the leaf, for example by using a distance field (Fig 5B). The growth directions are then projected onto this developmental axis, and separated into components that are parallel (Fig. 5D) and perpendicular (Fig 5E) to the axis. In Fig 5D a gradient of growth along the proximal-distal axis can be seen, with an increase in lateral growth around the forming serration (Fig 5E). This is not immediately apparent in the original PDG visualization (Figure 5C). Another benefit of looking at PDGs in the context of a local coordinate system is that it can provide a more direct comparison to the outputs of computational simulations. Developmental models of emergent organ shape often use morphogens that are thought to specifically control growth in the different directions in relation to a developmental axis (Kierzkowski et al., 2019; Kuchen et al., 2012; Whitewoods et al., 2020). By projecting the PDGs onto this axis, it is possibly to directly compare model growth rates in the different directions to experiments. Since MorphoGraphX can load a wide variety of mesh formats, this allows the direct comparison of similar quantifications made on templates extracted from model simulations from various sources.

**Figure 5:**
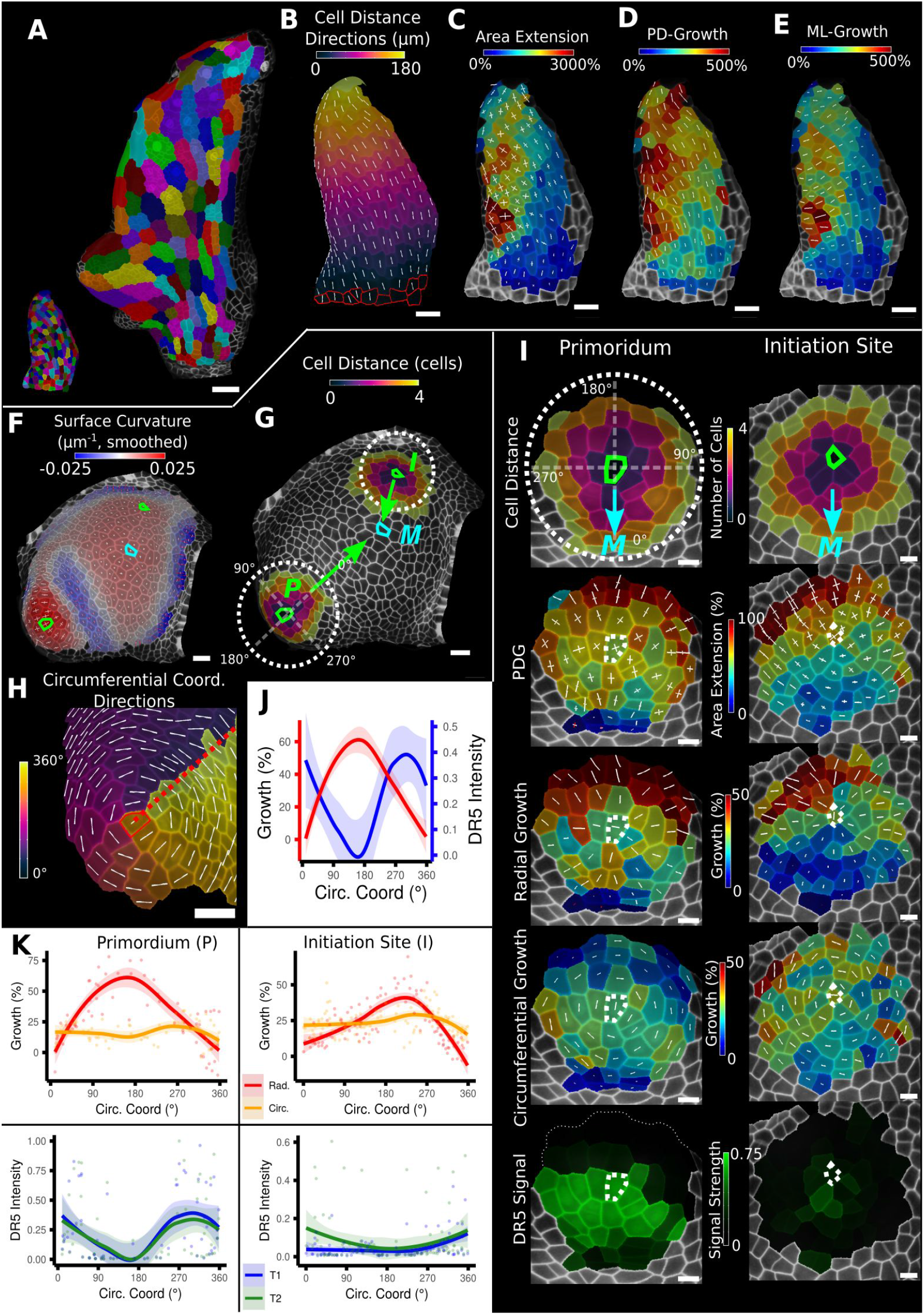
Examples of data analyses using organ coordinate directions. (A-E) Quantification of cellular growth along organ axes in a young *A. thaliana* leaf. (A) Segmented meshes of the leaf primordium at 3 and 6 days after initiation shown with cell labels and lineages of the earlier time point (3 days). (B) Earlier time point of (A) with proximal-distal (PD) axis coordinates (heat map) and directions (white lines) computed from selected cells at the leaf base. (C) Area extension (heat map) and Principal Directions of Growth (PDGs, white lines) between the time points of (A). PDG axes are computed per cell and can point in different directions. (D-E) Computation of the growth component of (C) that is directed along the PD and the orthogonal medial-lateral (ML) axis. (F-K) Quantification of locally directed growth in leaf primordium and initiation site of a tomato meristem. (F) Smoothed heat map of cell curvature. Local maxima in this heat map (green & cyan cells) were selected as meristem center (M), primordium center (P) and initiation site (I) as shown in (G). (H) To analyze the data we defined a circumferential coordinate system with its axes directions (white lines) around the primordium and initiation center (not shown), and aligned them towards the meristem center. (I) Heat maps of cell distance, area extension, radial and circumferential growth and normalized DR5 signal intensity of the aligned primordium and initiation site. (J) Plotting the data of (I) reveals a negative correlation of the DR5 signal intensity and radial growth around the developing primordium. (K) Detailed plots of radial (red) and circumferential growth (orange) as well as the normalized DR5 signal intensity of the primordium and initiation site. Scale bars: (A) 50 μm; (B - H) 20 μm; (I) 10 μm.

Figure 5F-K shows a similar growth quantification for the tomato meristem, where suitable local organ coordinates were created using cell distance measures around each emerging leaf primordium with directions pointing towards (radial) and around (circumferential) their respective center. Additionally, the signal intensity of the auxin reporter DR5 was quantified in the same sample. For both primordia we found radial growth to have a high negative correlation with DR5 signal intensity, whereas circumferential growth was more or less constant. The DR5 maximum tends to be on the adaxial side of the initiating leaf, whereas growth is much higher on the opposing abaxial side. This suggests that auxin acts as a trigger for primordium initiation, rather than via controlling growth rates directly.

The mature ovule in Arabidopsis shows a more complicated structure than a root or sepal, with four layers of integument cells encapsulating the nucellus that contains the embryo sac (Schneitz et al., 1995; Vijayan et al., 2021, Fig1S1, Fig 4). After segmentation and 3D mesh extraction in MorphoGraphX, directions normal to the surface (Fig. 4A) were combined with a Bezier curve computed from a user-selected cell file, and used to construct a coordinate system for the outermost cell layer (Fig. 4B,C). The Bezier curve defined the longitudinal direction whereas the surface directions obtained from the organ surface mesh were used to compute perpendicular width and depth axes and distances. Cell volume and geometry acquired from 3D segmentation and mesh extraction (Fig 1S1) were calculated along the various directions of the organ axes and analyzed (Fig. 4D-G).

Measuring the length, width and depth of cells along the cell layer and surface directions revealed the underlying cause for differences in cell volume between different proximal-distal regions of the outermost integuement layer. Moving along the proximal-distal axis, we found variations in cell volume with a clear minimum at around 100 μm and a steady increase towards the distal end (Fig. 4G). Cell length at the proximal and distal ends showed substantial differences, not seen in cell width and cell depth. The increase in cell length at the distal end was accompanied by enhanced cell anisotropy. These findings suggest that the overall steady rise in cell volume at the distal end is mainly due to differential cell length (Fig. 4G).

### Using positional information for automatic cell type classification

Plant organs typically emerge as primordia consisting of undifferentiated tissue. Cells subsequently differentiate, acquiring a unique identity that depends on their location within the organ, via genetic processes that integrate spatial and environmental cues. Although cell differentiation is ultimately controlled by differential gene expression, it is often the case that cell fate can be predicted by geometry, even at very early stages (Yoshida et al., 2014). It is rare that cells with different cell type have identical morphological features. Although most workflows in MorphoGraphX begin by segmenting 3D image stacks into cells, this is primarily an initial step to enable further analysis. As such, recent method development for MorphoGraphX has largely focused on the downstream processing of segmented data. The software now supports a large variety of measures to quantify different features of cell morphology, including: simple geometric measures (area, volume, perimeter, surface area, min and max axis), shape quantifiers (convexity, circularity, lobeyness, largest empty circle, aspect ratio), neighborhood measures (number of neighbors, variability), gene expression (average, total, near boundary), and cell network measures (betweenness centrality, betweenness current flow). Most measures can be used on time-lapse data to quantify changes over time (growth rates, gene expression changes, cell proliferation). For a complete list of the measures implemented in MorphoGraphX, see Supplemental tables 1, 2. The modular architecture of MorphoGraphX also allows custom measures tailored to specific problems to be easily added through its plug-in interface. More sophisticated calculations, for example the averaging of data over multiple samples, can be calculated externally in packages such as R and imported back into MorphoGraphX for visualization on segmented meshes. Here, the development of more complex data-flows is enabled by the use of a standardized attribute system to store and visualize cell data, for both scalar values and tensor (directional) information.

During the segmentation process MorphoGraphX assigns a unique label to each cell. A secondary cell label is provided (parent label), which is often used for lineage tracking. Other secondary labelings are also possible, for example cell type, cell layer, or zones within an organ. MorphoGraphX has several methods to assign these labels to classify cell types and layers. These labels can be assigned manually by interactively selecting cells, or by employing a number of processes that use heat map measure data to assign secondary labels (Fig. 3E-H). Positional information from the distance measures can be combined with measures of cell morphology and gene expression, where a secondary labeling can be used to provide additional context.

Since cells of a common biological cell type have similarities in one or more geometrical, positional or gene expression attributes, the values of these attributes will often form a cluster, facilitating their automatic classification. An example can be seen in the 3D Cell Atlas addon for MorphoGraphX (Fig. 3S1 A-C; T. D. Montenegro-Johnson et al., 2015) that clusters cells by relative radial distance and cell size to classify the cell layers of the root, hypocotyl, mature embryo, or other radially symmetric plant organs. To aid in optimizing cell clustering, MorphoGraphX offers a two dimensional interactive heat map, where information from two independent measures can be visualized, and clusters selected (Figure 3S1 B, 3G). These methods can be used repeatedly on sub-sets of cells to enable a classification of cells that differ across more than 2 features.

Multi-feature classifications tasks can be solved automatically by machine learning approaches (Cortes & Vapnik, 1995) when provided with sufficient training data. Of particular relevance are Support Vector Machines (SVMs) which have been used to classify cell types based on geometrical features of plant cells (de Reuille & Ragni, 2017; Sankar et al., 2014). MorphoGraphX provides a simple interface to the libSVM support vector machines library (Chang & Lin, 2011). Cells can be selected and classified into different cell types for use as training data (Fig. 3H, Movie 2). Any cell attribute or measure that can be quantified in MorphoGraphX can be used by the classifier. These include all the morphological and gene expression measures, time-lapse measures, the positional information created via distance maps or other coordinate systems, and even custom measures created via plugins or calculated externally with R or MATLAB. Once trained on a small group of cells with the desired measures, the classifier can be used to classify all the cells in a sample (Fig. 3H). After manual curation, the classification can then be used as additional training data, improving the power of the classifier. Figure 3G,H and 3S1 F-G shows the cell types of the Arabidopsis gynoecium, with stomata homogeneously distributed within the valve, consistent with the uniform growth and differentiation of this tissue (Eldridge et al., 2016; Ripoll et al., 2019).

**Figure 6S1:**
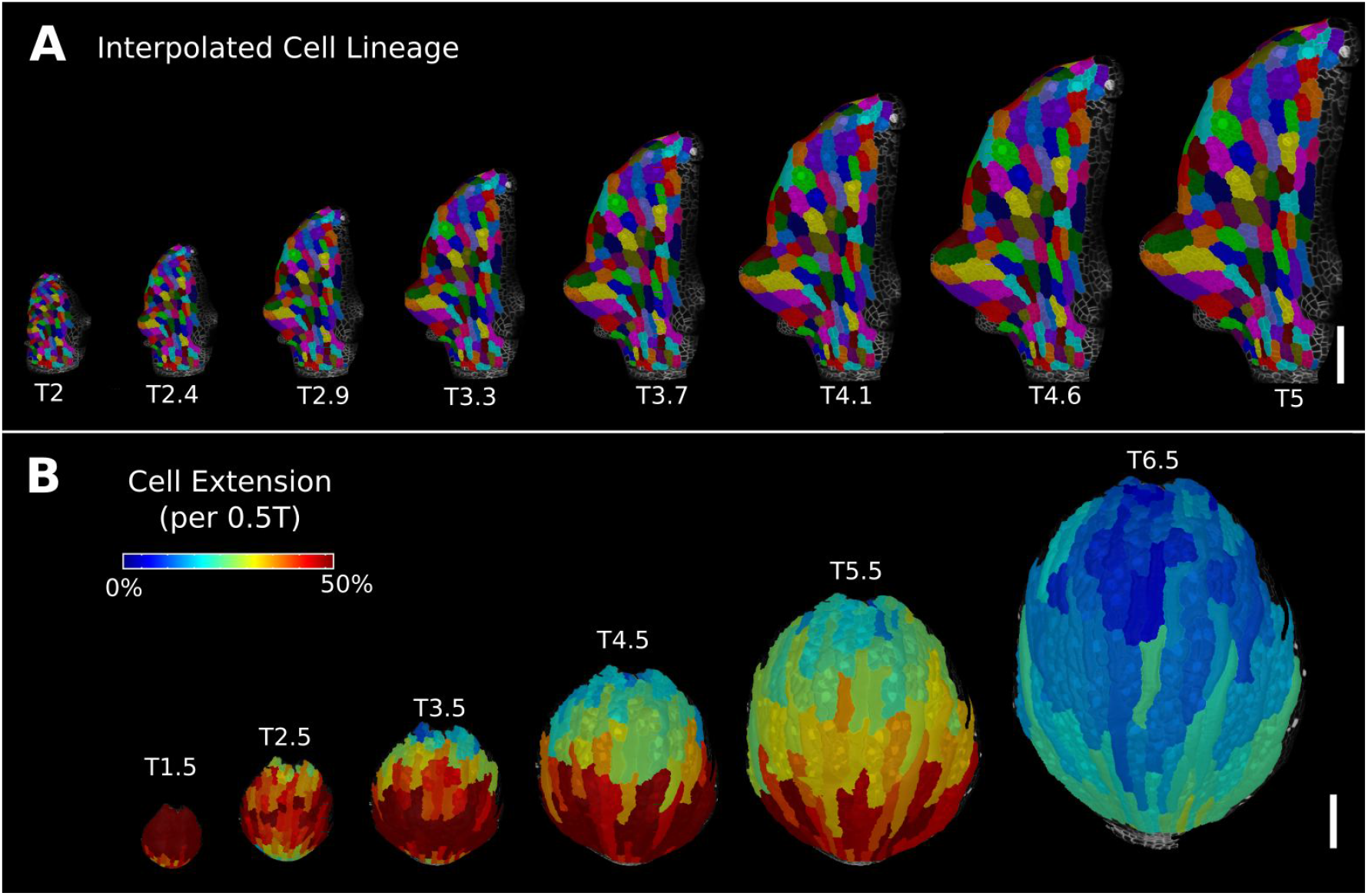
Deformation functions allow the interpolation of intermediate steps which can be turned into a continuous sequence or animation. (A) Animation of the early leaf development of *A. thaliana* created from T2 and T5 of Figure 5, shown with the lineages of T2. (B) Intermediate points of the animation of the sepal growth of *A. thaliana*. For the actual timepoints see Figure 2S1. Scale bars: 100μm.

### Mapping positional information through time

In the analysis of morphogenesis, many key quantifications such as growth depend on the ability to track samples through time. In MorphoGraphX this can be done following cell segmentation by manually assigning parent labels to the second time point, a process that has been highly streamlined in the user interface for 2.5D surfaces. However for full 3D samples, or large 2.5D samples with multiple time points, this method can be cumbersome. One method to address this problem is to find a non-linear coordinate transformation or deformation that maps all the points from one time point onto the next. Parent labeling can then be determined by mapping cell centers of the later time point to an earlier one, allowing the cell they came from to be identified. This can be used to directly assign lineage, or to seed algorithms that use more involved methods, such as the minimization of the total distances between the mapped cells (Fernandez et al., 2010). In MorphoGraphX, to define a mapping between the meshes of two successive time points (Fig 6A) we have implemented a 3D deformation function based on scattered data point interpolation (using cubic radial-basis functions; Duchon, 1977; Turk & O’Brien, 1999). An initial transformation is computed based on a few preassigned landmarks, by matching several cells with their parents in the previous time point (Fig 6C). A deformation field is then calculated which provides a mapping for all points in 3D. This is then used to assign parent labels in the second time point based on their closest match in the previous time point. Close to the landmark points, this mapping will be very accurate, with accuracy decreasing with distance from the landmarks. The decrease in accuracy away from landmarks is larger if the deformation between the time points is highly non-uniform. After assigning all the cells their closest parent, the mapping is then verified by checking the correspondence of neighborhoods between each cell and its parent. The labeling for cells which do not match is cleared, and the process is repeated. This causes correctly labeled regions to “grow” out from the initially placed landmarks, until the entire sample is correctly labeled (Fig 6C, Movie 3). At each step, only correctly labeled cells remain. Sometimes the iterative cell-labeling process can get stuck in highly proliferative areas where cells have divided repeatedly between time points. In this case a few additional landmarks can be manually added at trouble spots. One significant advantage of the method is that incorrect cells remain unlabeled, making manual curation straightforward. Once all of the parents are assigned and have passed the neighborhood correspondence check, one can be assured that both the lineage and the underlying segmentations are correct.

**Figure 6:**
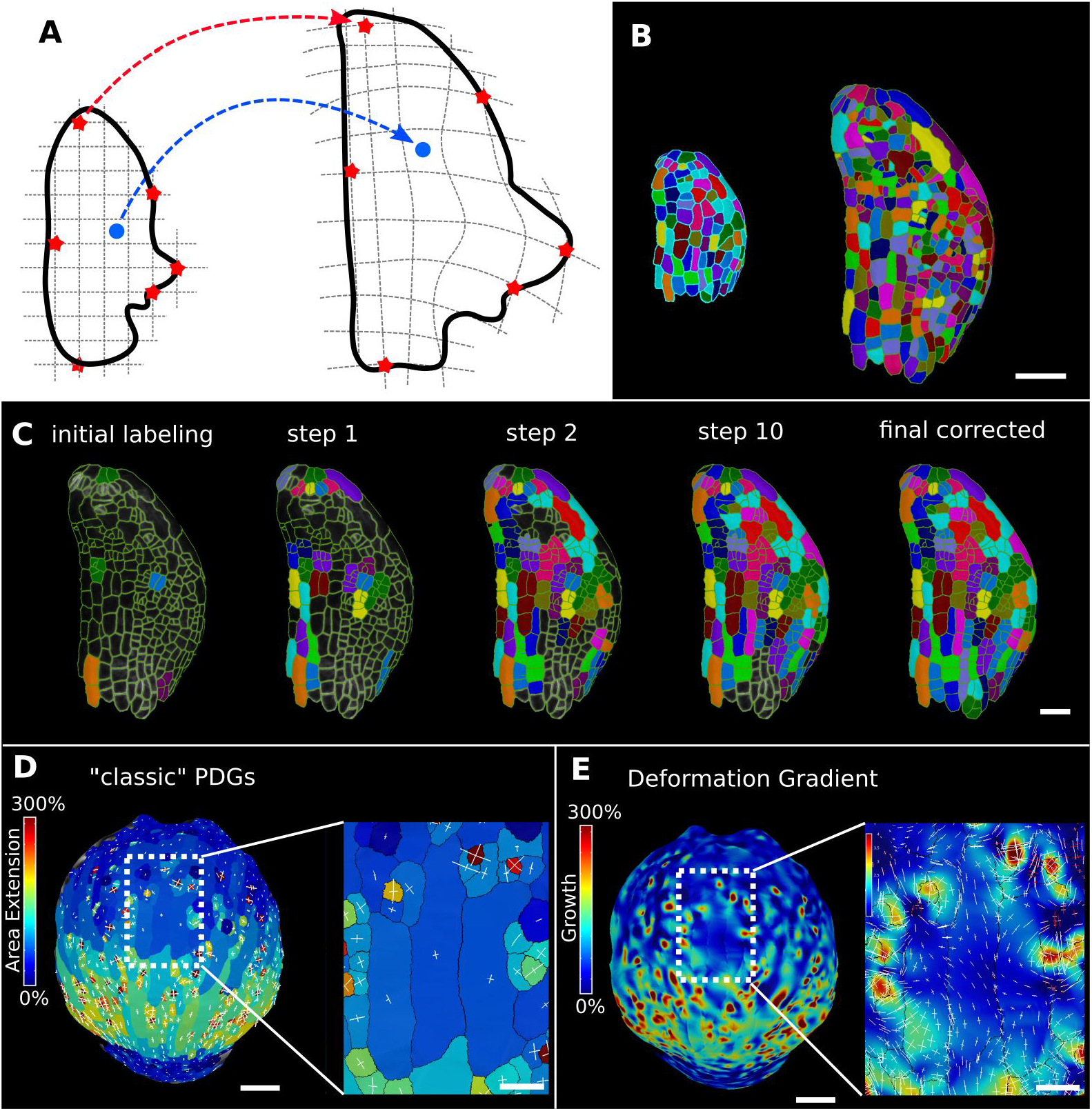
Deformation functions in MorphoGraphX. (A) Deformation functions allow a direct mapping of arbitrary points (blue) between two meshes. They require the definition of common landmarks (red stars). (B-C) Semi-automatic parent labelling using deformation functions. (B) Two consecutive time points of an *A. thaliana* leaf primordium segmented into cells. (C) The automatic parent labelling function requires the definition of a few manually labelled cells as initial landmarks. From this sparse correspondence, a mapping between the meshes can be created and new cell associations between the two meshes are added and checked for plausibility. With more cells found, the mapping between the meshes is improved for the next iteration. (D-E) Comparison of the classic Principal Directions of Growth (PDGs) in (D) with the gradient of a deformation function computed using the cell junctions from a complete cell lineage in (E) on an *A. thaliana* sepal. The classic PDGs compute a deformation for each cell individually and are shown with a heat map of areal extension for each cell. In contrast, the deformation function is a continuous function on the entire mesh. Here heat values are derived by multiplying the amount of max and min growth. Using the deformation function gradient reveals subcellular growth patterns that were previously hidden, such as differential growth within a single giant cell. Scale bars: (B, D, E) 50μm; (C & zoomed regions in D & E) 20μm.

Deformation functions can also be used to create animations of organ development from 2.5D or 3D time-lapse data. This requires two or more time points of a segmented mesh with corresponding cell lineages. The cell centers and/or junctions are used as the landmarks defining the deformation function that maps one mesh onto the other. Interpolating mesh vertices between stages creates a smooth animation with as many intermediary steps as desired (Fig. 6S1, Movie 4). MorphoGraphX has a user-friendly pipeline to record animations directly from the GUI with options to adjust the camera angle and to visualize cell lineages, heat maps and cell outlines during the animation. Temporal smoothing of morphing animations created from more than two time points is achieved using Catmull-Rom splines to interpolate the position of mesh vertices over time (Catmull & Rom, 1974). Heat and signal values in the mesh, such as cell area, growth rates or gene expression can also be interpolated along with vertex positions.

In large cells, growth can vary significantly within the same cell (Armour et al., 2015; Elsner et al., 2018). As the deformation function provides a smooth mapping between time-points, its gradient can be used to create a continuous growth map at any point on a mesh. This enables the approximation of areal expansion and PDGs at a sub-cellular level, where the quality of the approximation is limited by the number and placement of landmarks (junctions). It is also possible to apply the process to subcellular landmarks, such as those obtained by tracking microbeads, as done previously for 2D images (Armour et al., 2015; Elsner et al., 2018). Our 3D implementation of this method has been used to compute growth directions on curved surface meshes (Fig 6E) and volumetric meshes (Fig 7I).

**Figure 7:**
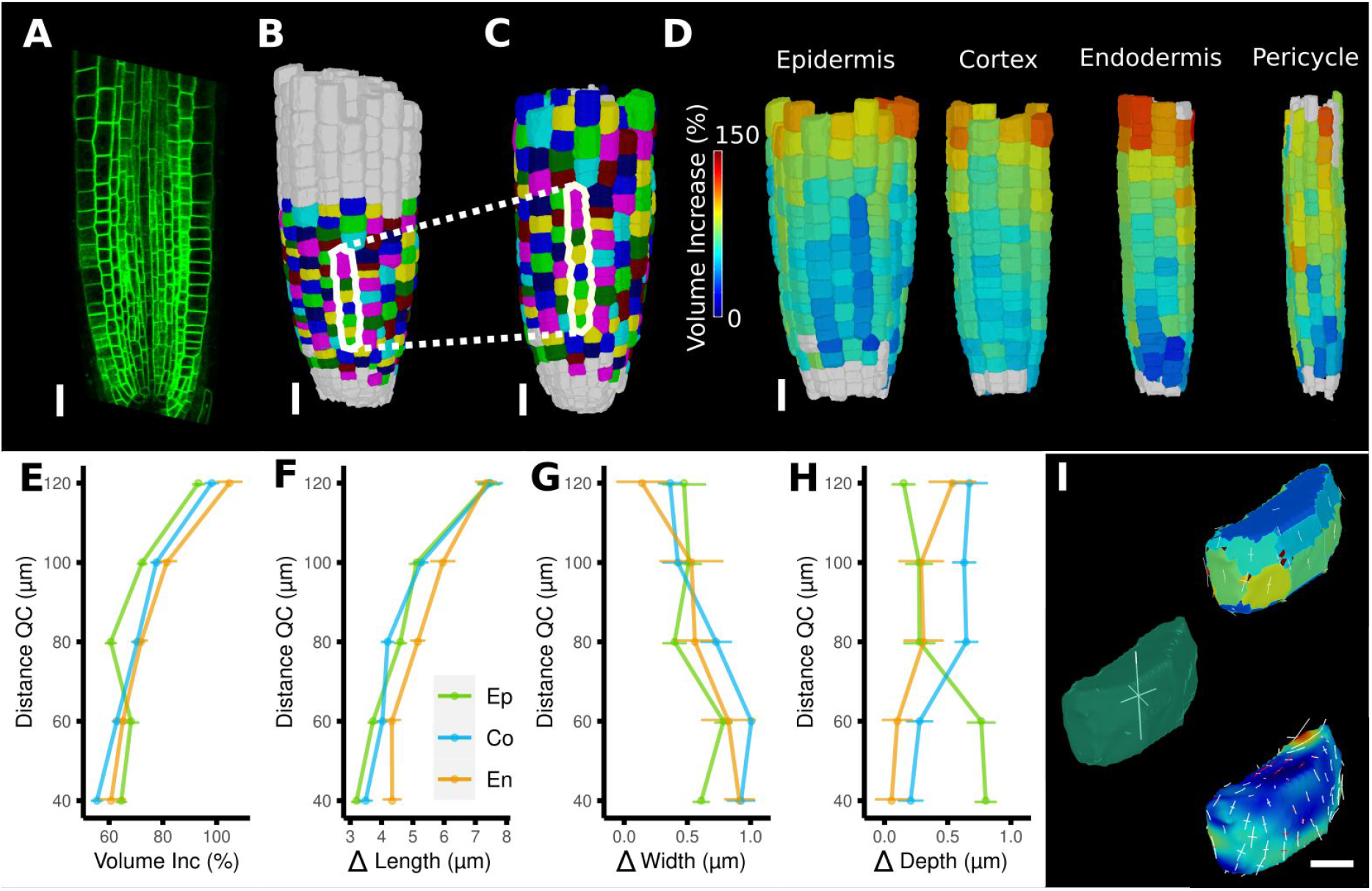
Time-lapse analysis and visualization of 3D meshes. (A) Cross section of the confocal stack of the first time point of a live imaged *A. thaliana* root. (B-C) The 3D segmentations of two time points imaged 6 hours appart. Shown are the cell lineages which were generated using the semi-automatic procedure following a manual correction. (D) Exploded view of the second timepoint with cells separated by cell types (see also Figure 4D). Cells are heat colored by their volume increase between the two time points. (E-H) Quantification of cellular growth along different directions within the organ. (E) Plot of the heat map data of (D). The cellular data was binned based on the distance of cells from the QC. Shown are mean values and standard deviations per bin. (F-H) Similarly binned data plots of the change in cell length (F), width (G) and depth (H). It can be seen that the majority of growth results from an increase in cell length. See Figure 7S1 for a detailed analysis of the cells in the endodermis. (I) Different ways to visualize 3D growth demonstrated using a single cortex cell: PDGs averaged over the cell volume (left), PDGs averaged over the cell walls projected onto the walls (top right), subcellular vertex level PDGs projected onto the cell walls (bottom right). Scale bars: (A-D) 20 μm; (I) 5 μm.

A comparison of the areal growth and PDGs calculated with deformation functions vs the cell-based method is shown in Figure 6D, E. The deformation function captures differences in growth within single cells, as is often apparent in larger cells that straddle areas of fast and slow growth. Figure 7A-I shows a 3D time-lapse of the Arabidopsis root where the deformation functions have been used to perform lineage tracking in 3D. It can be seen in Fig 7D that the 4 tissue types, epidermis, cortex, endodermis and pericyle all show a similar growth pattern, with slow growth in the meristem and faster growth near the transition zone. When the growth is displayed as a function of the distance from the root tip (Fig 7 E-H), again it can be seen that the layers are almost the same, reflecting the almost completely anisotropic growth of this system.

**Figure 7S1:**
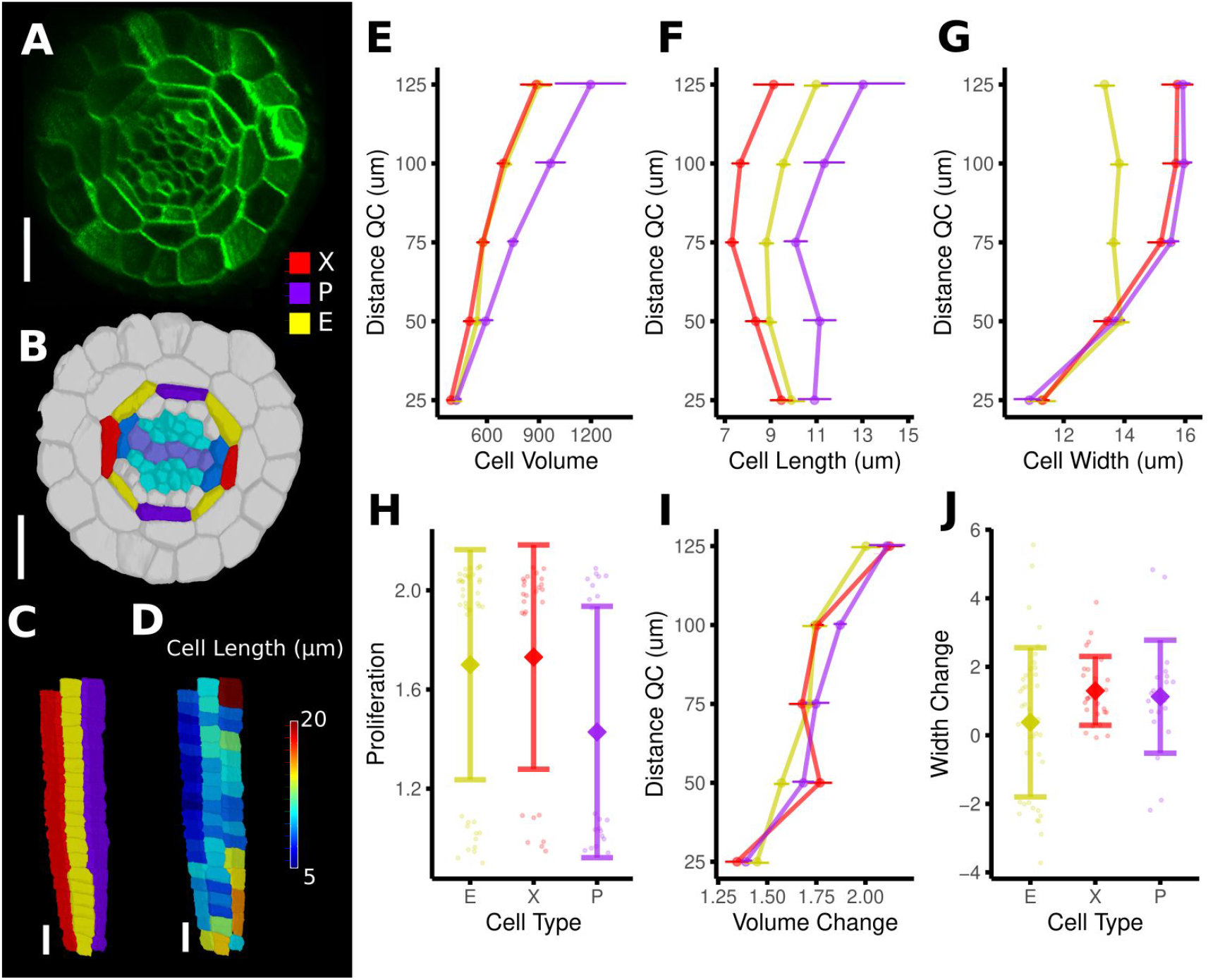
Time-lapse analysis of cellular geometry in the *A. thaliana* root endodermis. (A) Cross section of the confocal image of time point 1. (B) Segmentation with extended cell type labelling in the endodermis. The root cell type labelling of Fig. 3D was extended by identifying the xylem cells (light purple) in the stele (cyan), their adjacent pericycle cells (blue) and assigned the endodermis cells neighboring those pericycle cells as xylem file cells (X, red). Then the endodermis cells at right angles to the xylem files were assigned phloem file (P, purple) and the remaining endodermis cells (E, yellow). (C) Side view of one cell file of each cell type. As the cell types do not change along the cell file, it was possible to automatically assign the cell files based on their circumferential coordinate. (D) Cell files of (G) with a heat map of cell length indicating smaller cells in the xylem pole. (E-J) Quantifications of cell geometry and development in the endodermis cell types. Cellular data was binned according to their distance from the QC (E,F,G,I). Shown are mean values and standard deviations per bin (E,F,G,I) or cell type (H,J). P cells showed a larger volume (E), which was caused by a greater cell length (F), an observation which has been made before by (Andersen et al., 2018). In contrast, X and E cells were smaller in volume due to different reasons: While X cells were the shortest (E), E cells showed a lower cell width with increasing distance from the QC (G). The time-lapse analysis confirmed above observations: While volume change was similar across the cell type (I), P cells showed a lower proliferation rate (H), whereas E cells showed the smallest extension of cell width (J).Scale bars: (A, B) 20 μm; (C, D) 50 μm.

### Advanced geometric analysis

While MorphoGraphX was created to work with 2.5D surface projections, it now supports a complete set of tools for full 3D image processing, and in many samples advantages can be gained from combining both techniques. Additional tools, such as the automated cell lineage tracking on surfaces, have also been extended to 3D to facilitate growth analysis in full 3D. This is a much harder task than the analysis of surface images, as 3D cellular meshes lack the relatively easy to identify junctions that serve as material points for surfaces. However in many cases entire organs are well defined by their surfaces meshes, allowing landmarks on the surface to be used to construct a 3D deformation function to aid lineage tracking in full 3D. Surface landmarks can also be combined with 3D cell centers and/or the face centers as material points to improve the internal resolution of the deformation functions for full 3D samples. These techniques allow the methods used to calculate growth rates and PDGs in 2.5D to be extended to full 3D (Fig. 7I). Cell proliferation and most of the other measures can also be quantified from 3D time-lapse data (Fig 7S1). In addition to the automated tools, improved manual 3D parent labeling and the ability to relabel cells so that adjacent cells are always a different color, aid in the manual curation of 3D lineage maps.

One of the more advanced quantifications from time-lapse data is the analysis of cell division. As plant cells cannot move, cell division and growth are the main determinants of morphogenesis. MorphoGraphX has processes to identify dividing cells from time-lapse data, and quantify the orientation of the division wall in both 2.5D and 3D (Fig 8A-I, Fig. 8S1). In 2.5D the best fit line to the division wall is calculated (Fig 8A,D), whereas in 3D the best fit plane is used (Fig 8B,C,G). There are also measures to determine the asymmetry of the daughter cells. The use of positional information to give organ context is even more important for directional information. For instance, quantifying the orientation of the division plane is of little use without knowing how it relates to the developmental axes. If the organ shape is simple and aligned with one of the axes, then cell division orientations can be used directly (Fig 8S1A). When more complex shapes are involved, orientations can be computed with respect to the axes of a local coordinate system defined for the organ, along with its associated positional information (Fig. 8E,I). It is also possible to quantify how close cell divisions are to common division rules proposed in the literature, such as the shortest wall through the center of the cell including local minima (Fig 8H; Besson & Dumais, 2011), along the principal directions of growth (Hejnowicz, 1984), or rules based on network measures (Jackson et al., 2019).

**Figure 8:**
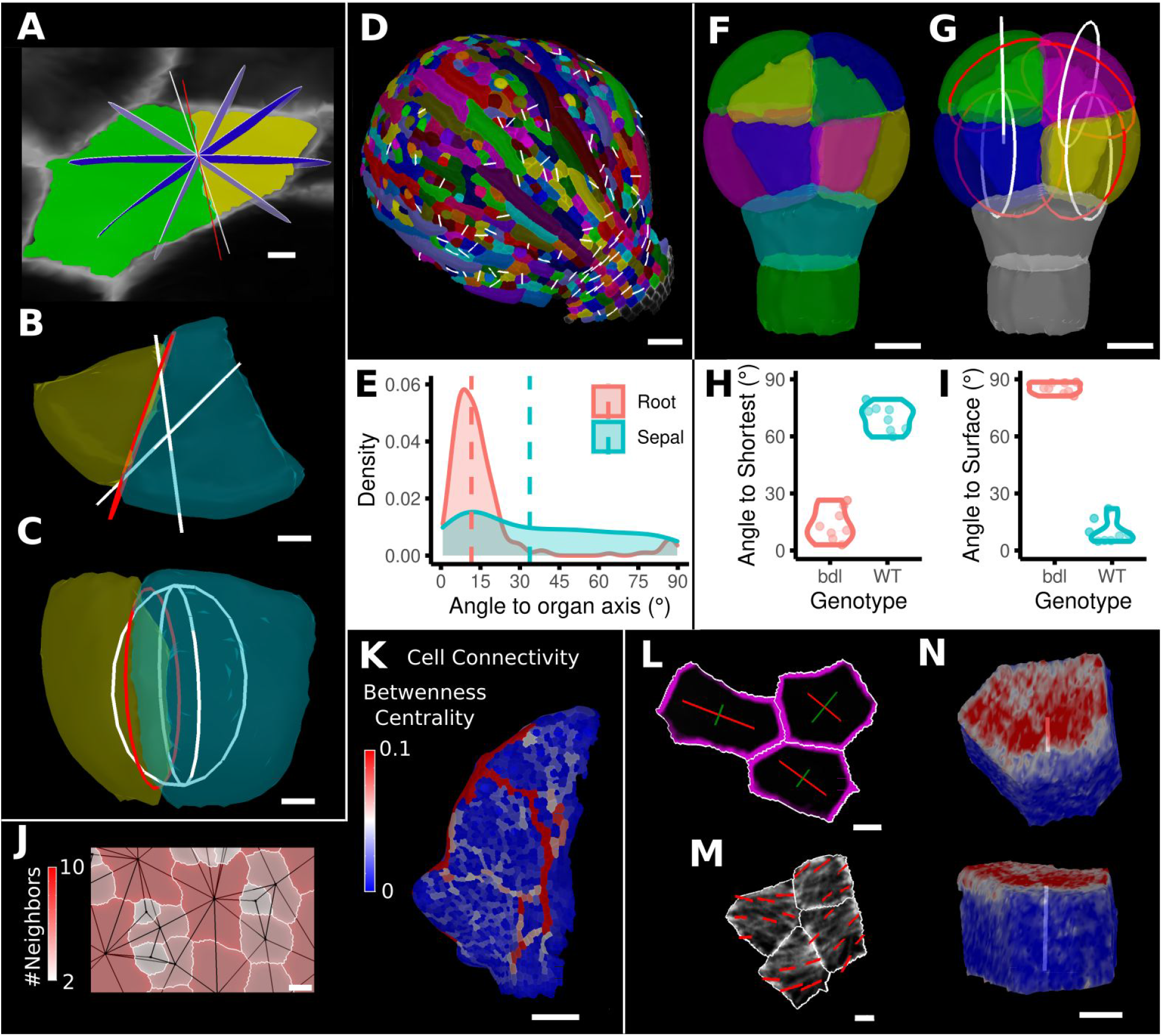
Advanced data analysis and visualization tools. (A) Division analysis of a cell from a surface segmentationl of an *A. thaliana* sepal. A planar approximation of the actual plane is shown in red and other potential division planes in white/blue. The actual wall is very close to the globally shortest plane. (B-C) Top and side view of a recently divided 3D segmented cell. The daughter cells are colored yellow and cyan. The red circle depicts the flat approximation plane of the actual division wall. The two white rings depict the two smallest area division planes found by simulating divisions through the cell centroid of the mother cell (i.e. the combined daughter cells). (D) Visualization of the actual planes (white lines) between cells that divided into 2 daughter cells in the *A. thaliana* sepal. (E) Density distribution and median (dashed line) of the angle between the division plane and the primary organ axis in sepal (see D) and root (see Figure 8S1A). The division in sepals are less aligned with the organ axis. (F) Half of an *A. thaliana* wild type embryo in the 16 cell stage. This view shows that the divisions leading to this stage are precisely regulated to form 2 distinct layers in the embryo. (G) A visualization of the actual planes (red circles) and the shortest planes (white circles) in the wild type. Cells are colored according to the label of the mother cells. (H-I) Violin plots of quantifications of the planes show that the wild type does not follow the shortest wall rule, unlike the auxin insensitive inducible bdl line RPS5A>>bdl. The bdl divisions are almost orthogonal to the organ surface (see Figure 8S1 B,D,E), whereas the wild type divides parallel to the surface. Consequently, the bdl fails to form a distinct inner layer. (J-K) Cellular connectivity network analysis. (J) Cell connectivity network analysis on a young *A.thaliana* leaf. Cells are heat colored based on the number of neighbors. (L) Heat map of betweenness centrality. The betweenness reveals pathways which might be of importance for informtaion flow, potentially via the transport of auxin. (L-N) Cell based signal analysis. (L) Analysis of cell polarization on a surface mesh. (M) Microtubule signal analysis on a surface mesh. (N) Top and side view of a cell polarization analysis on a volumetric mesh (root epidermis PIN2, see Figure 8S2A-D for details). Scale bars: (A,B,C,L,M) 2 μm; (D) 50 μm; (F,G,J,N) 5 μm; (K) 100 μm.

The organization of cells in organs may be analyzed through the extraction of cell connectivity networks from 2.5D or 3D segmented data. The physical associations between cells (cell-cell wall areas) can be extracted and converted into networks where they are analyzed using network measures (Fig. 8J-K). Local measures such as the number of immediate neighbors (degree) can be calculated, along with more global measures, such as betweenness centrality based on the number of shortest paths cells lie upon, or random walk centrality (Fig. 8K). These global measures are central to understanding how information flows within tissues (Jackson et al., 2017, 2019). The use of these measures uncovered the presence of a global property in cellular organization within the Arabidopsis SAM (Jackson et al., 2019). Namely, the length of paths between cells is maximized, whereby cells which lie upon more shortest paths have a great propensity to divide, and the orientation of this division tends to leave the two daughter cells on the fewest number of shortest paths. Using this approach, the local geometric properties of cells can be related to the emergent global organization of cellular arrangements. Perturbation of cell shape in the *katanin1* mutant led to alterations in path length in the SAM, which correlated with defects in phyllotaxis (Jackson et al., 2019).

Reporter signals, such as proteins tagged with fluorescent proteins, can also be quantified in MorphoGraphX. After segmenting a surface into cells by using a cell wall stain or marker line, a signal collected in a second channel can be projected onto the surface mesh, and the abundance, orientation and polarity of signals can be computed. Examples are the PIN-FORMED (PIN) auxin transporter report line (Benková et al., 2003) or the GFP:MBD (Van Bruaene et al., 2004) line that tags microtubules (MTs), (Figure 8L-N, 8S2). For the quantification of cell polarity the projected signal along the cell border is binned based on its position in relation to the the cell center to obtain its predominant direction and its intensity. Figure 8L shows an example of the PIN1 polarity quantification at the cell wall of surface segmented cells in the SAM. A similar quantification can be performed for 3D meshes as shown in Figure 8N and Figure 8S2A,B where we computed the PIN2 polarity in epidermis and cortex cells of an Arabidopsis root. Again, this directional information can be combined with the organ coordinates to compute the angle between cell polarity and the organ axis (Figure 8S2C). Another example is the quantification of MT alignment using an implementation of Fibril tool (Boudaoud et al., 2014) that has been adapted for processing on surfaces. After projecting the MT signal onto the surface, the alignment direction and strength of the signal can be quantified at the sub-cellular level (Figure 8M) or for entire cells (Figure 8S2E). Again, this information can be interpreted using organ coordinates, as we demonstrate on cells of a SAM which tend to have their MTs aligned circumferentially from the meristem center (Figure 8S2E).

**Figure 8S1:**
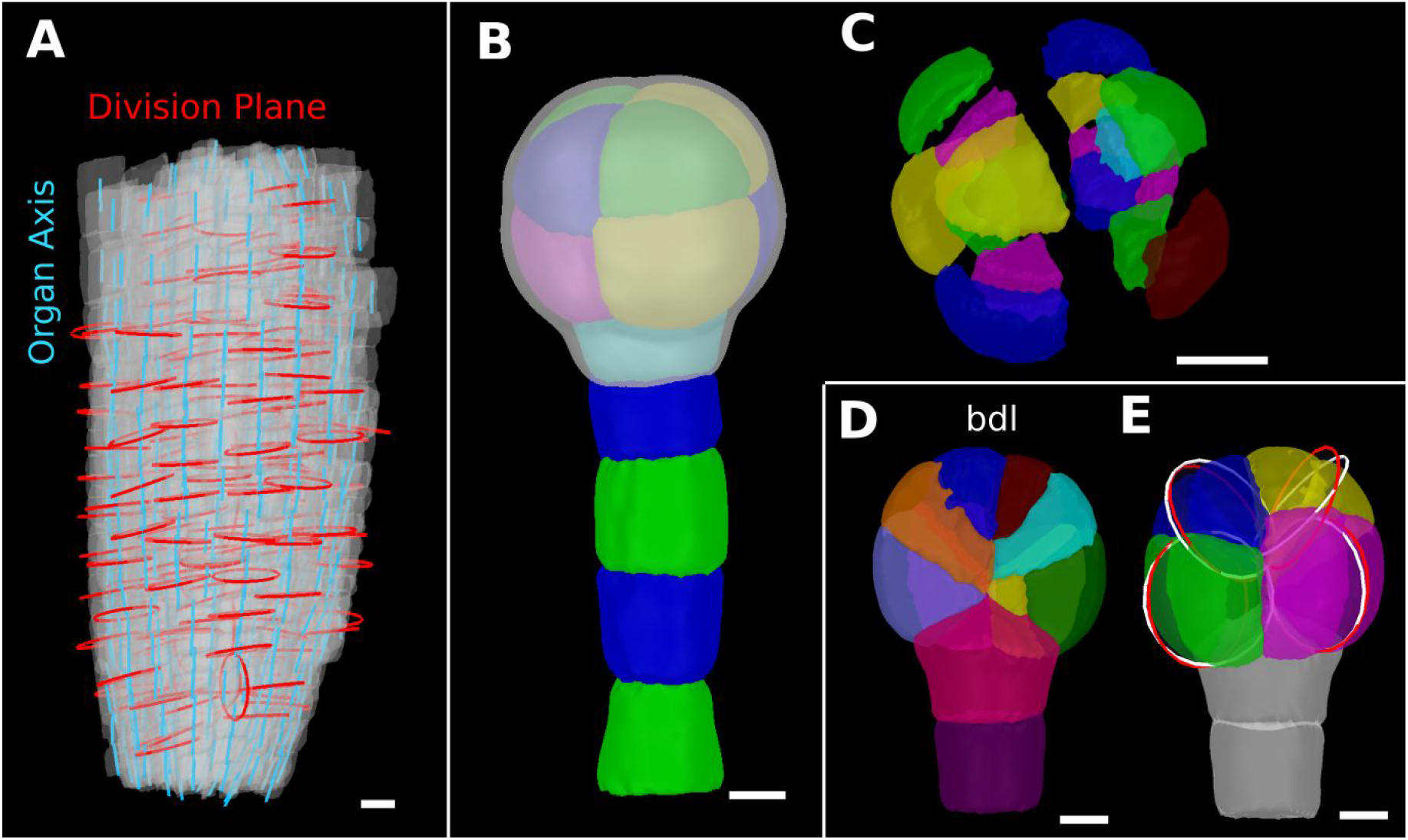
Details of the cell division analysis examples from Figure 8. (A) Second time point of an *A. thaliana* root (see Figure 7C) that was used for the division plane analysis in Figure 8E. Cells are shown semi-transparently (grey) with their longitudinal organ axis (cyan) obtained from the analysis using 3D Cell Atlas in Figure 4B. Planar approximations of the division planes between cells that divided between the 2 time points are shown as red circles. Consistent with the quantitative analysis in Figure 8E most planes are aligned with the organ axis. (B-C) Wild type embryo at the 16-cell stage segmented into volumetric cells shown with an organ surface mesh (grey, semi-transparent, B) and shown in an exploded view (C) to enable the visualization and access of inner layers. (D-E) Corresponding panels to Figure 8F-G for bdl embryo. Scale bars: (A,B,C) 10 μm; (D,E) 5 μm.

**Figure 8S2:**
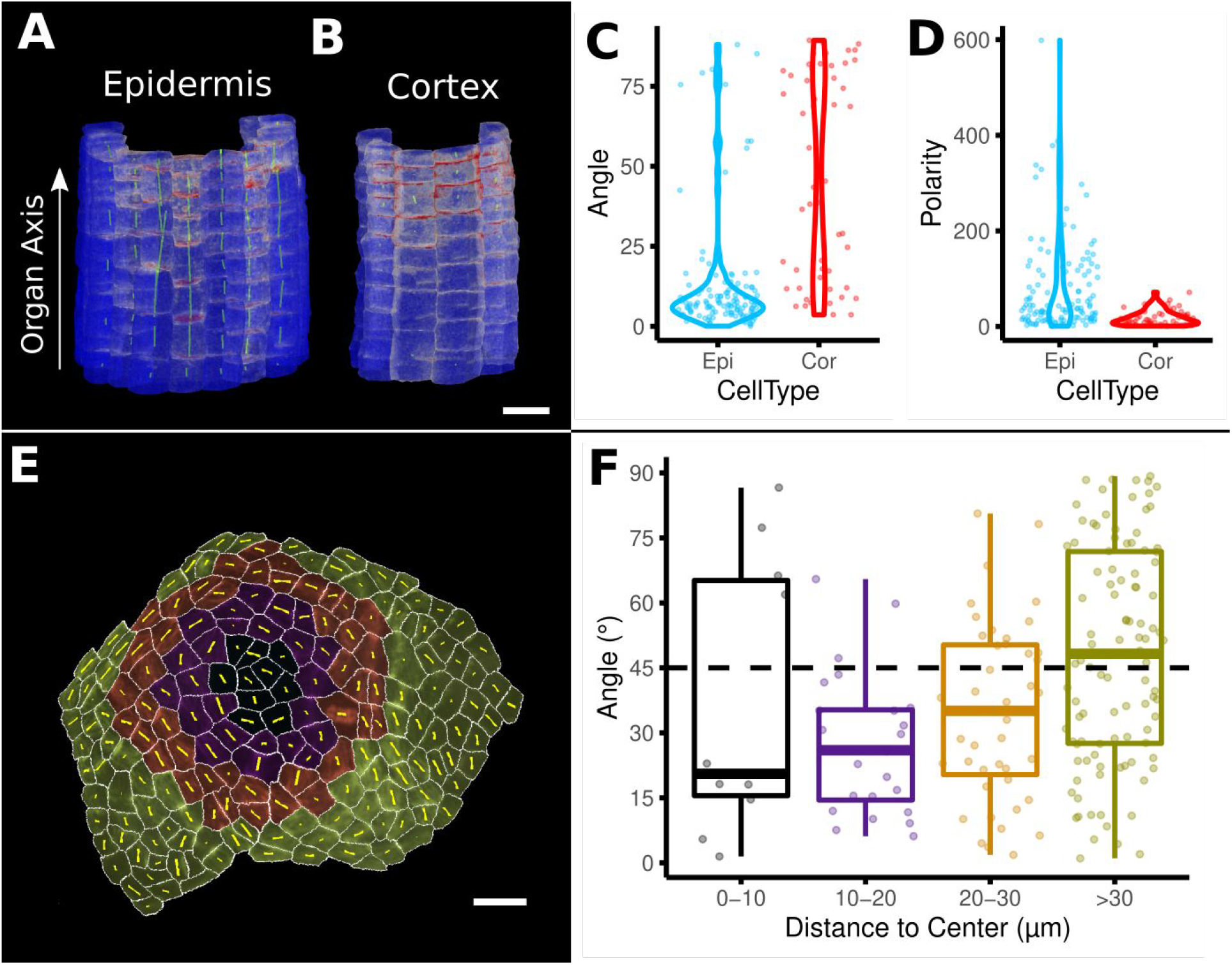
Example analyses of cell polarity and microtubule signals of the data shown in Figure 8M-N. (A-D) Quantification of PIN2 polarity in volumetric cells. (A-B) Heat map of PIN2 concentration on epidermis and cortex of an *A. thaliana* root. Green lines depict directionality and strength of the PIN2 concentration. (C-D) Violin plots of the orientation data for division planes and PIN2 polarity for epidermis and cortex cells. Epidermis cells show considerably stronger polarity (D) and are more aligned with the (longitudinal) organ axis (C). (E-F) Microtubule analysis on a SAM of *A. thaliana*. (E) The cells on the SAM were binned according to their distance to the SAM center. Cells are heat colored according to their bin. Yellow lines show the direction and strength of the microtubule orientation. (F) Boxplot of the angular difference between microtubule orientation and the circumferential direction around the center of the SAM (similar to Figure 5H). Scale bars: (A,B) 20 μm; (E) 10 μm.

### 3D visualization and interactive tools

MorphoGraphX has a flexible rendering engine that can handle meshes containing millions of triangles. It supports the independent rotation and translation of different stacks and meshes in the same world space and the ability to render both voxel and geometric data together with blending and transparency. It has adjustable clipping plane pairs and a bendable Bezier cutting surface that can be used to look inside 3D samples, and an interface to support the creation of animations (Movie 5). However, visualizing and interacting with 3D data on a 2D computer screen still remains a challenge. A particular problem is the validation and correction of 3D segmentations of organs, as internal cells are obscured by outer layers. Exploded views are a commonly used method to visualize the internal structure of multi-component 3D objects (Fig 8S1 C; Li et al., 2008), which can be used on mesh data in MorphoGraphX. To add biological meaning to the exploded visualization, cells can be bundled by their parent or cell type label to visualize key aspects of biological data sets such as cell divisions or to separate organs into cell layers or by cell type (Fig 7D, 8S1C). These processes can also aid user interactions, making cell selection and annotation more straightforward when users are curating 3D cellular segmentations and lineage maps.

In addition to mesh editing tools, MorphoGraphX has several tools to edit voxel data. The simplest is an eraser tool which can be used to remove portions of the stack that would otherwise interfere with processing. An example is the digital deletion of the peripodial membrane overlying the Drosophila wing disc, which needs to be removed to allow for the extraction of the organ’s surface (Aegerter-Wilmsen et al., 2012). A typical workflow for 3D segmentation starts with a 3D image of a cell boundary marker. This is then pre-processed with operations such as blurring to reduce noise or background removal filters, before segmentation with algorithms such as the ITK (www.itk.org) morphological watershed. More recently, deep learning methods with convolutional neural networks (CNN) have been developed to predict cell boundaries, such as the 3D U-Net model (Çiçek et al., 2016), that can improve the stacks for downstream segmentation. The modular structure of MorphoGraphX has allowed us to interface an implementation of the 3D U-Net model developed by Eschweiler et al. (2019). This enables the interactive use of the CNN boundary prediction tool from within MorphoGraphX (Movie 6), and simplifies experimentation with different networks or downstream segmentation strategies. It also avoids the requirement to set up a full Python environment, and the associated installation issues. Although some tools have packaged 3D U-Net prediction with a choice of several segmentation algorithms (Wolny et al., 2020), the tools are typically written as pipelines without intermediate visualization of the data, making it cumbersome to experiment with the different methods and parameters of the individual data processing steps in the pipeline.

Once the data is segmented, it often requires some manual correction before it is ready for final analysis in a chosen coordinate system. MorphoGraphX has interactive tools that operate on voxel data both to combine and split labels (cells), although typically it is easier to over-segment and combine, rather than under-segment and split (Movie 6). This can be used to correct segmentations, which can then be used to help train deep learning networks to further improve automatic segmentation (Vijayan et al., 2021). In this context MorphoGraphX has been used to segment and curate Arabidopsis ovule data to create ground truth for confocal prediction networks (Wolny et al., 2020).

## Conclusion

Similar to sequencing data, geometric data on the shape and sizes of hundreds or thousands of cells is of limited value without annotation. For many developmental questions, the spatial context for information on cell shape, division and gene expression is paramount. However it is not enough to know the 3D position of cells, but rather their position in a coordinate system relative to the developing tissue or organism. These coordinates would typically reflect the developmental axes of the organism or tissue, allowing the direct comparison of cell and organ shape changes with the gene expression controlling their morphology. In addition to putting data in a mechanistic context, organ-centric relative coordinates can be used to compare samples with different morphologies (Thompson, 1942), such as different mutants, or even in different species (Kierzkowski et al., 2019). This also applies to changes in morphology over time, where organ coordinate systems can be used to determine the correspondence of cells at different stages of development. Several tools have been used to successfully harness organ-centric coordinates for specific problems (Montenegro-Johnson et al., 2015, 2019; Schmidt et al., 2014), and MorphoGraphX provides a generalized framework to make coordinate systems customized to the particular organ or organism of interest.

In addition to annotation with organ-centric positional information, MorphoGraphX has a comprehensive toolkit for cell shape analysis, growth tracking, cell division, and the quantification of polarity markers, both on 2.5D image meshes, and for full 3D. All of these measures can be calculated and stored within the mesh, or exported to files for further processing with other software. Custom measures can also be calculated and imported for visualization within MorphoGraphX. Cell shape measures in combination with positional information provide a powerful framework for cell type classification, both with machine learning methods (Fig 3G-H) as well as clustering techniques (Montenegro-Johnson et al., 2019).

A key strength of MorphoGraphX is that it offers both manual and automatic tools for segmentation, lineage tracking and data analysis. Although fully automated methods are improving, streamlined methods for manual and semi-automatic segmentation and analysis provide a path to completion for many samples where the automatic methods are “almost” good enough. For example, the automatic lineage tracking now available benefits from the streamlined tools we developed previously to do the process manually, as these are now used to correct and fill in missing portions when the automatic segmentation is incomplete. This reflects the interactive nature of MorphoGraphX and its focus on low-throughput but high-quality data sets. Nevertheless, portions of a workflow can be fully automated by using python scripts to process many files. Furthermore, operations performed in MorphoGraphX are written to a python log file, allowing easy cut-paste script creation.

The internal architecture of MorphoGraphX has been designed to make it easily extendable, while retaining the speed of the fully compiled, statically typed language C++. The relatively small visualization and data management core is augmented with processes that are loaded dynamically at startup and provide almost all of the software’s functionality. MorphoGraphX has grown to provide a wealth of custom processes for 2.5D image processing and coordinate system creation. Additionally, it has become a platform to integrate published tools and methods that have no visual interface to the data of their own, which increases their accessibility and ease of exploration for biologists. Examples include the previously mentioned CNN process, and the interface to the XPIWIT software (Bartschat et al., 2016) which provides a tool to develop ITK image processing pipelines interactively without any C++ programming required. These pipelines can be packaged into plugins and called directly from MorphoGraphX, allowing the exploration of different ITK filters. One XPIWIT pipeline we have pre-packaged in a process is for the TWANG method (Stegmaier et al., 2014), a fast parallel algorithm for nuclear segmentation. Another example of software integration is the processes we have developed to interface with R (Movie 7) that provides plots for basic statistical analysis on attributes created in MorphoGraphX, including positional information provided by organ-centric coordinate systems. This simplifies the creation of the most commonly used plots without the need for export files.

As more and more imaging datasets are becoming available for community use, their annotation with positional information and gene expression data will be critical to help understand how the cell-level action of different genes and genetic networks is translated into the 3D forms of tissues and organs of different species. In this context MorphoGraphX provides a tool set to help maximize the attainable information from these datasets, in an accessible platform tailored to the experimental biologist.

## Material and Methods

### Software Availability

MorphoGraphX is available for Linux and Windows, although we recommend Linux as some add-ons are only available for Linux, and some, such at the CNN add-on require Cuda. For Linux we provide a Cuda version for machines with a compatible nVidia graphics card, and a non-Cuda version for those without. Currently there is only a non-Cuda version for Windows. Although there is no Mac version, some have had success running it in a virtual machine. The software and documentation are available at www.MorphoGraphX.org.

### Data Acquisition

#### Arabidopsis Flower meristem (Fig1)

pUBQ10::acyl:TDT (Segonzac et al., 2012) and DR5v2::n3eGFP (Liao et al., 2015) were crossed. F3 double homozygote line was used for imaging. Floral meristems were dissected from 2 weeks old plants grown on soil under the long-day condition (16 h light/ 8h dark), at 20°C ± 2°C using injection needle. Dissected samples were cultivated in 1/2 Murashige and Skoog medium with 1% sucrose supplemented with 0.1% plant protective medium under the long-day condition (16 h light/ 8h dark), at 20°C ± 2°C. Confocal imaging was performed with Zeiss LSM800 with a 40× long-distance water immersion objective (1 NA, Apochromat). Excitation was performed using a diode laser with 488 nm for GFP and 561 nm for TDT. Signal was collected at 500-550 and 600-660 nm, respectively. Images of 3 replicates were obtained every 24 hours for 4 days.

#### Arabidopsis Ovule (Fig1S1, Fig4A-F)

Data previously published in (Vijayan et al., 2021).

#### Arabidopsis Root (Fig2A, Fig 3B,D, Fig3S1A,C, Fig7A-D,I, Fig7S1A-D, Fig8S1A)

Root imaging: pUBIQ10::H2B-RFP pUBQ10::YFP-PIP1;4 was described previously (von Wangenheim et al. 2016). The seeds were stratified for 1 day at 4°C, grown on 1/2 Murashige and Skoog medium with 1% sucrose under the long-day condition (16 h light/ 8h dark) at 20°C ± 2°C. Confocal imaging was performed with Zeiss LSM780 with two-photon laser (excitation 960nm) with a band pass filter 500/550nm for YFP. Images of 3 replicates were obtained.

#### Arabidopsis Mature Embryo (Fig2C)

Arabidopsis thaliana Col-0 seeds were sterilized in 70% ethanol with Tween20 for two minutes, replaced with 95% ethanol for 1 minute and left until dry. Seeds were placed on the Petri plates containing 1/2 MS medium including vitamins (at pH 5.6) with 1.5% agar and stratified at 4°C for 3 days in darkness. Next, seeds were imbibed for 3 hours, and the mature seed embryo was isolated from the seed coat. For live imaging, the embryos were stained with propidium iodide 0.1% (Sigma-Aldrich) for 3 minutes and imaged with Leica SP8 laser scanning confocal microscope with a water immersion objective (x20). Excitation wavelengths and emission windows were 535 nm and 617 nm. Confocal stacks were acquired at 1024×1024 resolution, with 0.5-μm distance in Z-dimension. Images were acquired at 48 hours intervals and samples were kept in a growth chamber under long-day condition (22°C, with 16 h of light per day) between imaging. From more than 10 replicates a sample with curved overall shape was selected for the demonstration of the organ coordinates using a Bezier curve. To quantify the cell area, change and anisotropy, the fluorescence signal was segmented, and cells were parent labeled manually between two successive time points. Heat-maps are displayed on the later time-point (after 48 hours of growth). Scale bars are displayed on the image.

#### Marchantia time-lapse (Movie 3)

Marchantia polymorpha gemmaling Cam-1 PM::GFP reporter line (Shani et al., 2010) were transferred on a Petri plate containing 1/2 Gamborg’s B5 medium including vitamins (pH 5.5) with 1.2% agar and grown for 24 hours. For live imaging, the gemmaling were imaged with Leica SP5 laser scanning confocal microscope with a water immersion objective (x25/0.95). Excitation wavelengths and emission windows were 488 nm and 510 nm. Confocal stacks were acquired at 1024×1024 resolution, with 0.5-μm distance in Z-dimension. Images were acquired at 24 hours intervals and samples were kept in a growth chamber under constant light between imaging. For the move we selected a representative sample from 6 total replicates. To quantify the cell area, change and anisotropy, the fluorescence signal was segmented and semi-automated parent labelling as performed to couple the cells at two successive time points. Heat-maps are displayed on the later time-point (after 24 hours of growth). Scale bars are displayed on the image.

#### Arabidopsis Sepal (Fig2D,E, Fig2S1A,B, Fig6D,E, Fig6S1B, Fig8A,D)

Data previously published in (Hervieux et al., 2016).

#### Arabidopsis Leaf (Fig5A-E, Fig6S1A, Fig8J,K)

Data previously published in (Kierzkowski et al., 2019).

#### Tomato SAM (Fig5F-I)

Data previously published in (Kierzkowski et al., 2012).

#### Arabidopsis SAM (Fig3A,C, Fig3S1D)

Data previously published in (Montenegro-Johnson et al., 2019).

#### Arabidopsis Gynoecium (Fig3E-H) and Leaf (Fig 6B,C)

pUBQ10::acyl:YFP has been described previously (Willis et al., 2016). Plants were cultivated on soil under the long-day condition (16 h light/ 8h dark), and 20°C ± 2°C. Flowers at post-anthesis stage from 5 weeks-old plants were dissected with fine tweezers to remove sepals and stamens to expose gynoecium and mounted on the 60 mm plastic dish filled with 1.5% agar. Confocal imaging was performed with a Zeiss LSM800 upright confocal microscope, equipped with a long working-distance water immersion objective 40X (1 NA, Apochromat). Excitation was performed using a diode laser with 488 nm for YFP and the signal was collected between 500 and 600 nm. For both organs images of 3 replicates each were obtained.

#### Arabidopsis Embryo (Fig8B,C,F,G, Fig8S1B-E)

Data previously published in (Yoshida et al., 2014).

#### Arabidopsis Meristems for PIN1 and MT (Fig8L,M, Fig8S2A-C)

pUBQ10::acyl:TDT (Segonzac et al., 2012) and GFP:MBD (Van Bruaene et al., 2004) were crossed. F3 double homozygote line was used for imaging.

Floral organs were removed with fine tweezers about 21 days after germination to expose inflorescence meristem. Meristems were mounted on the 60 mm plastic dish filled with 1.5% agar and imaged with a Zeiss LSM800 upright confocal microscope, equipped with a long working-distance water immersion objective 60X (1 NA, Apochromat). Excitation was performed using a diode laser with 488 nm for GFP and 561 nm for TDT. Signal was collected at 500-550 and 600-660 nm, respectively. Images of 3 replicates were obtained.

#### Arabidopsis Root for PIN2 in 3D (Fig8N-O, Fig8S2E)

pPIN2::PIN2:GFP was previously described (Xu & Scheres, 2005). The seeds were stratified for 2 days at 4°C, grown on 1/2 Murashige and Skoog medium with 1% sucrose under the long-day condition (16 h light/ 8h dark) at 20C° ± 2°C. The roots were stained by 10mM propidium iodide (Sigma-Aldrich), and observed by Zeiss LSM780 with two-photon laser (excitation 990nm) with a band pass filter 500/550nm for GFP and 575-600nm for PI. Images of 3 replicates were obtained.

### Data Analysis

For the data analysis examples in the paper we computed all necessary cellular data within MorphoGraphX and exported them as csv files. Those files were imported to RStudio for further processing or directly plotted using ggplot2 (R Core Team, 2020; RStudio Team, 2020; Wickham, 2016).

#### Arabidopsis Flower meristem (Fig. 1)

We selected one sample for segmentation and further analysis. The segmentation, cell lineages and heat maps were generated following the standard workflow as described in (Strauss et al., 2019) and in the MGX user guide.

#### Arabidopsis Ovule (Figures 1S1, Fig. 4)

Segmentation was obtained by back blending the raw images to CNN boundary predictions (Wolny et al., 2020) as described in (Vijayan et al., 2021).

For the analysis in Fig. 4 we selected one sample of the published data.The trimmed surface mesh was used to extract the outermost layer using the method described in (Montenegro-Johnson et al., 2019). The organ axis was defined by a Bezier curve obtained from a manually selected central cell file using the processes “Misc/Bezier/Bezier From Cell File” and “Mesh/Cell Axis 3D/Custom/Create Bezier Grid Directions”. Moreover, for each cell the direction towards the surface and the orthogonal direction of the Bezier and surface direction was computed (“Create Surface Direction”; “Create Orthogonal Direction”), resulting in 3 orthogonal organ axes. The cell sizes were quantified by first doing a PCA on the voxels of cells on the segmented stack (“Mesh/Cell Axis 3D/Shape Analysis/Compute Shape Analysis 3D”) and finally computing the component of the PCA’s tensor aligned with the axes of interest (“Mesh/Cell Axis 3D/Shape Analysis/Display Shape Axis 3D” with the appropriate “Custom” heat option), with the Bezier direction corresponding to cell length, the surface direction to depth and the orthogonal direction to width.

Shape Anisotropy was defined using the equation: (max - 0.5*min - 0.5*min)/(max+mid+min), with max, mid and min defined by the length of the PCA axes.

To create the plots, cells with a distance >40μm from the central cell file were ignored. The data of the remaining 213 cells was plotted.

#### Arabidopsis Root (Figures 2A, 4B,D, 4S1A,C, 7A-D,I, 7S1A-D, 8S1A)

From the 3 replicates we selected the sample with the best segmentation quality for further analysis. The two time points of the analyzed root data sample were segmented using ITK segmentation processes in MGX (see also (Stamm et al., 2017) and the MGX user guide). From the segmented stack a surface mesh and volumetric cell mesh were obtained. For the axis alignment analysis (Fig 2A), the organ was manually aligned with the y-axis and the coordinates of the cell centroids were computed. In Fig 2B the cell volume of the 304 epidermis cells was plotted. The 3D Cell Atlas pipeline (Montenegro-Johnson et al., 2015; Stamm et al., 2017) was used to compute cell coordinates, sizes and cell types (Fig 4B, D; Fig4S1A-C).

For the time-lapse analysis (Fig 7A-I, Fig 7S1) the cell lineages were determined semi-automatically using the pipeline introduced in this paper (Fig 6) followed by a manual error correction. PDGs in 3D were derived from the deformation function mapping the first onto the second time point using parent labelled cell centroids and cell wall centers.

For the analysis of the cell types in the endodermis (Fig 7S1), xylem cells in the stele and their neighboring pericycle cells were automatically identified by their circumferential coordinate. Endodermis cells touching two xylem-associated pericycle cells were determined as xylem pole. The two phloem poles in the endodermis were shifted by 90 degrees (or 2 cells) from the xylem pole. In total, we used 26 xylem pole, 21 phloem pole and 43 rest endodermis cells for the analysis.

For the analysis in Fig 8E and Fig 8S1A only the second time point was used. We computed the proliferation to the previous time point, extracted the vertices on each division plane between cells that have divided exactly once (proliferation = 2, n=249 mother cells that divided) and computed a PCA on each set of division plane vertices to extract the normals of the planes. Then we computed the angle between the longitudinal axis of the organ as extracted by 3D Cell Atlas and the division planes and exported to data.

#### Arabidopsis Sepal (Figures 2D,E, 2S1A,B, 6D,E, 6S1B, 8A,D)

For the sepal analysis one replicate of the data from (Hervieux et al., 2016) consisting of 7 time points was used (see Fig 2S1A,B).

For the analysis in Fig 2D-F and Fig2S1 for each time point we manually determined the organ base based on the cell lineages from the first time point. Cells at the organ base were selected and used to compute the Euclidean cell distance measure. Finally, cell distances, growth, proliferation and cell sizes were exported.

For the cell division analysis in Fig 8A,B we analyzed the divisions that occurred between the time point T4 and T5 (n=84). We computed the proliferation between these time points, extracted the vertices on each division plane between cells that have divided exactly once (proliferation = 2) and computed a PCA on each set of division plane vertices to extract the normals of the planes. Next, we computed the PD-axis direction of the organ using the Euclidean cell distance from the base. Finally we computed the angle between PD-axis and the division planes and exported to data.

For the growth analysis in Fig 6D we computed the PDGs from time point T4 to time point T5, visualized on the earlier time point. Fig 6E shows the same time point, but here growth was computed using the gradient of the deformation function obtained from the cells’ junctions.

#### Arabidopsis Shoot Apical Meristem (Figures 3A,C, 3S1D)

We selected one sample of the published data for analysis. Cell type labels were determined using the methods described in 3D Cell Atlas Meristem (Montenegro-Johnson et al., 2019).

#### Arabidopsis Leaf (Figures 5A-E, 6S1A, 8J-K)

The Arabidopsis leaf data was previously published in (Kierzkowski et al., 2019). One replicate of a time-lapse series consisting of 7 time points was selected for analysis, but only time points T2 and T5 were used. The cell distance was computed similar to the Sepal example (Fig 2D) as distance from the organ base. Additionally we computed the heat map gradient of the cell distance heat map (“Mesh/Cell Axis/Custom/Create Heatmap Directions”) to obtain custom direction along the proximal distal (PD) axis and orthogonal to that the medial-lateral (ML) axis of the organ for each cell. PDGs were computed and used to determine the amount of growth along the previously computed PD and ML-axis.

For the cell network analysis in Fig 8J-K we computed the cell connectivity network of all cells in T5 weighted by the inverse of the length of the cell walls to determine the betweenness centrality (“Mesh/Heat Map/Measures/Network/Betweenness Centrality”), (Jackson et al., 2019).

#### Tomato Shoot Apical Meristem (Figure 5F-I)

For the growth and DR5 signal analysis on the shoot apical meristem we used one replicate of the previously published data of (Kierzkowski et al., 2012). To objectively find the center of the meristem, the primoridium and the initiation site the curvature of the cells was computed. The resulting heat map was smoothed across neighboring cells for two rounds and resulting local maxima were identified as centers. Meshes were manually aligned along the x-axis with respect to the meristem center to compute circumferential coordinates (“Mesh/Heat Map/Measures/Location/Polar Coord”) around primordium and initiation center. For the analysis only cells in the vicinity of the primordium and initiation centers were considered. These cells were determined using a threshold in the cell distance measure from those centers. Furthermore, the gradients of the Euclidean cell distance heat maps from both centers were used to compute custom directions along the heat (=radial) and orthogonal to the heat (=circumferential). Now the growth analysis was done similar to the leaf with computing PDGs and growth along the custom axis.

#### Arabidopsis Shoot Apical Meristems (Figures 8L,M, 8S2E,F)

For the MT analysis we selected one sample for segmentation and analysis. We determined the center of organ based on a smoothed curvature heat map. The center cell was selected and the Euclidean cell distance to the remaining cells was computed. The circumferential direction around the cell center was obtained from the orthogonal direction of the heat map directions. Cells were then binned by their Euclidean distance to the center.

#### Arabidopsis Embryo (Figures 8F,G, 8S1B-E)

The data for the 3D division analysis in *A. thaliana* embryos was previously published in (Yoshida et al., 2014). From this data set we chose one wild-type and one inducible bdl (pRPS5a>>bdl) sample at the 16 cell stage.

A surface mesh was generated from the cells in the embryo and the cells were parent labelled according to their predicted mother cell. Then the process “Mesh/Division Analysis/Analysis 3D/Division Analysis Multi” performed the following steps on all of the parent labelled cells (n=16 cells or 8 divisions in each genotype): First a planar approximation of the actual division plane was computed by performing a PCA on the vertex positions of the shared wall between the two daughter cells. Then 1000 equally distributed division planes were simulated on the combined mother cell and different measures were quantified. The actual and the best planes were visualized using “Mesh/Division Analysis/Display and Filter Planes”.

#### Arabidopsis Root PIN2 in 3D (Figures 8N, 8S2A,B)

For the analysis of the PIN directions in the *A. thaliana* root we selected one sample for segmentation. Next, we defined the main organ axis using a Bezier curve through the center of the organ. Then we computed the PIN2 polarity direction and determined the angle between the polarity direction and the Bezier line.

## Acknowledgments

We acknowledge support by the Center for Advanced Light Microscopy (CALM) of the TUM School of Life Sciences, and inspiration from the many collaborators we have worked with during the development of MorphoGraphX.

## MorphoGraphX 2.0 Supplemental Materials

**Supplemental Table 1:** Measures for cells on surface projections (2.5D).

| Measure | Unit | Description |
| --- | --- | --- |
| **Geometry** |  |  |
| Area | um<sup>2</sup> | Area of the Cell (sum of its triangle area) |
| Aspect Ratio | - | Ratio of Length Major Axis and Length Minor Axis (see below) |
| Average Radius | um | Average distance from the center of gravity of a cell to its border |
| Junction Distance | um | Max or min distance between neighboring junctions of a cell |
| Length Major Axis | um | Length of the major axis of the 2D Shape Analysis: (Computes a PCA on the triangle positions weighted by their area) |
| Length Minor Axis | um | Length of the minor axis of the 2D Shape Analysis: (Computes a PCA on the triangle positions weighted by their area) |
| Maximum Radius | um | Maximum distance from the cell center to its border |
| Minimum Radius | um | Minimum distance from the cell center to its border |
| Perimeter | um | Sum of the length of the border segments of a cell |
| **Lobeyness** |  |  |
| Circularity | - | $\text{Perimeter}^2 / (4 * \pi * \text{Area})$ |
| Lobeyness | - | Ratio of cell perimeter and convex hull perimeter. 1 for convex shapes |
| Rectangularity | - | Ratio of cell area and the area of the minimum rectangle that can contain the cell. 1 for rectangular shapes, lower values for irregular shapes. |
| Soldarity | - | Ratio of the convex hull area and the cell area. 1 for convex shapes, higher values for complicated shapes. |
| Visibility Pavement | - | Simply 1-Visibility(Stomata) |
| Visibility Stomata | - | Estimate of visibility in the cell. Returns 1 for convex shapes, and decreases with the complexity of the contour. |
| **Location** |  |  |
| Bezier Coord | um | Associated Bezier coordinate of a cell. Requires a Bezier grid. |
| Bezier Line Coord | um | Associated Bezier coordinate of a cell. Requires a Bezier curve. |
| Cell Coordinate | um | Cartesian coordinate of a cell |
| Cell Distance | um/<br>cells | Distance to the nearest selected cell (finds the shortest path to a selected cell, different distance measures: euclidean, cell number or 1/wall area) |
| Distance to Bezier | um | Euclidean distance to the Bezier curve or grid |
| Distance to Mesh | um | Euclidean distance to the nearest vertex in the other mesh |
| Major Axis Theta | ° | Angle between the long axis of the cell and a reference direction |
| Polar Coord | °/um | Polar coordinate around a specified Cartesian axis |
| **Network** |  |  |
| Neighbors | count | Number of neighbors of a cell |
| Betweenness Centrality | - | Computes the betweenness centrality of the cell connectivity graph. Edges can be weighted by the length of the shared wall between neighboring cells |
| Betweenness Current Flow | - | Computes the betweenness current flow of the cell connectivity graph. Edges can be weighted by the length of the shared wall between neighboring cells |
| **Signal** |  |  |
| Signal Border |  | Average amount of border signal in a cell |
| Signal Interior |  | Average amount of interior signal in a cell |
| Signal Parameters |  | Advanced and general process which allows the setting of parameters to compute different kinds of signal quantifications |
| Signal Total |  | Average amount of total (=border + interior) signal in a cell |
| **Other Measure processes** |  |  |
| Mesh/Lineage Tracking/Heat Map Proliferation | cells | Proliferation |
| Mesh/Cell Axis/Custom/Custom Direction Angle | ° | Angle between a Cell Axis and a Custom Axis |
| Mesh/Division Analysis/Compute Division Plane Angles | ° | Angle between division planes and/or cell axes |

**Supplemental Table 2:** Measures for meshes with volumetric (3D) cells

| **Measure** | **Unit** | **Description** |
| --- | --- | --- |
| **Cell Atlas** |  |  |
| Cell Length (Circumferential, Radial, Longitudinal) | um | Cell length as determined by 3D Cell Atlas Root (Shoot rays from the cell center to the side walls to measure cell size along organ-centric directions) |
| Coord (Circumferential, Radial, Longitudinal) |  | 3D organ coordinates as determined by 3D Cell Atlas Root |
| **Geometry** |  |  |
| Cell Length (Custom, X, Y, Z) | um | Cell length along the specified direction (cell size is measured as in 3D Cell Atlas Root (see above)) |
| Cell Wall Area | um<sup>2</sup> | Total area of the cell wall |
| Cell Volume | um<sup>3</sup> | Volume of the cell |
| Outside Wall Area | um<sup>2</sup> | Cell wall area that is not shared with another neighboring cell |
| Outside Wall Area Ratio | % | Proportion of Cell Wall Area that is not shared with a neighbor cell |
| **Location** |  |  |
| Bezier Coord | um | Associated Bezier coordinate of a cell. Requires a Bezier curve or grid. |
| Cell Coordinate | um | Cartesian coordinate of a cell |
| Cell Distance | um/cells | Distance to the nearest selected cell (finds the shortest path to a selected cell, different distance measures: euclidean, cell number or 1/wall area) |
| Distance to Bezier | um | Euclidean distance to the Bezier curve or grid |
| Mesh Distance | um | Euclidean distance to the nearest vertex in the other mesh |
| **Network** |  |  |
| Neighbors | count | Number of neighbors of a cell |
| Betweenness Centrality | - | Computes the betweenness centrality of the cell connectivity graph. Edges can be weighted by the length of the shared wall between neighboring cells |
| Betweenness Current Flow | - | Computes the betweenness current flow of the cell connectivity graph. Edges can be weighted by the length of the shared wall between neighboring cells |

